# Single-Mitochondrion Sequencing Uncovers Distinct Mutational Patterns and Heteroplasmy Landscape in Mouse Astrocytes and Neurons

**DOI:** 10.1101/2024.06.13.598906

**Authors:** Parnika S Kadam, Zijian Yang, Youtao Lu, Hua Zhu, Yasemin Atiyas, Nishal Shah, Stephen Fisher, Erik Nordgren, Junhyong Kim, David Issadore, James Eberwine

**Affiliations:** Department of Pharmacology, Perelman School of Medicine, University of Pennsylvania, Philadelphia, PA 19104, USA; Department of Bioengineering, School of Engineering and Applied Science, University of Pennsylvania, Philadelphia, PA 19104, USA; Department of Biology, School of Arts and Sciences; University of Pennsylvania, Philadelphia, PA 19104, USA

**Keywords:** single-mitochondrion, single-nucleotide variants, neurons, astrocytes, heteroplasmy

## Abstract

**Background:** Mitochondrial (mt) heteroplasmy can cause adverse biological consequences when deleterious mtDNA mutations accumulate disrupting ‘normal’ mt-driven processes and cellular functions. To investigate the heteroplasmy of such mtDNA changes we developed a moderate throughput mt isolation procedure to quantify the mt single-nucleotide variant (SNV) landscape in individual mouse neurons and astrocytes In this study we amplified mt-genomes from 1,645 single mitochondria (mts) isolated from mouse single astrocytes and neurons to 1. determine the distribution and proportion of mt-SNVs as well as mutation pattern in specific target regions across the mt-genome, 2. assess differences in mtDNA SNVs between neurons and astrocytes, and 3. Study cosegregation of variants in the mouse mtDNA.

**Results:** 1. The data show that specific sites of the mt-genome are permissive to SNV presentation while others appear to be under stringent purifying selection. Nested hierarchical analysis at the levels of mitochondrion, cell, and mouse reveals distinct patterns of inter- and intra-cellular variation for mt-SNVs at different sites. 2. Further, differences in the SNV incidence were observed between mouse neurons and astrocytes for two mt-SNV 9027:G>A and 9419:C>T showing variation in the mutational propensity between these cell types. Purifying selection was observed in neurons as shown by the Ka/Ks statistic, suggesting that neurons are under stronger evolutionary constraint as compared to astrocytes. 3. Intriguingly, these data show strong linkage between the SNV sites at nucleotide positions 9027 and 9461. f

**Conclusion:** This study suggests that segregation as well as clonal expansion of mt-SNVs is specific to individual genomic loci, which is important foundational data in understanding of heteroplasmy and disease thresholds for mutation of pathogenic variants.

## Background

Mitochondria are subcellular organelles that govern cellular processes including bioenergetics, calcium buffering, DNA repair, cell cycle, cell death, and organelle signal transduction (1). In addition to mitochondrial (mt) diseases (2), these mt-driven cellular processes are dysregulated in Parkinson’s (3), Alzheimer’s (4), cardiovascular and renal diseases (5, 6), and cancer (7). There can be as many as thousands of mitochondria in a cell depending on the cell type and metabolic state, and each mt can have up to tens of copies of circular, self-replicating mtDNA (8). The circular mtDNA in mice and humans is about 16.3 and 16.5 kb respectively and encodes 13 key proteins involved in the electron transport chain, and two rRNAs (namely 12S and 16S rRNA interspersed by 22 tRNAs (9, 10). Compared with the nuclear genome, mtDNA are at least 10 times more mutation prone (11), and both germline and somatic mutations can contribute to the coexistence of the wild-type and mutant alleles in the same organelle or the same cell, a phenomenon known as “heteroplasmy” (12). Pathogenic mt heteroplasmy is implicated in maladies including myoclonus epilepsy with ragged-red fibers (MERRF) (2) and Huntington’s disease (13), while pervasive mt heteroplasmy was also observed in cells of healthy humans and mice (11, 14–16).

It is known that mtDNA mutations accumulate over time and can reach a “disease-threshold” causing deleterious biological consequences. One approach to understanding this process is to study and decipher the patterns of SNV distribution and accumulation enriched by selective replication (17). To better understand the mechanistic aspects of such changes, the systematic structure of the mt heteroplasmy should be explored in context of its nested hierarchy i.e., the level of single mitochondria, single cells, and single organisms. Key questions such as heteroplasmy across different levels, the segregation mode of heteroplasmic alleles, and the level(s) of regulation upon which selection forces may exert, have been only partially addressed. Matthews *et al.* (18) showed that mitochondria segregated in heteroplasmic units in human fibroblasts, while studies by Cavelier *et al.* (19) reported predominantly homoplasmic nucleoids in human fibroblasts. Due to technical limitations, these early pioneering studies only queried single sequence loci in mtDNA. Evidence for tissue specific selection has been observed in mice (20) and more recently in humans (21), but these studies focused at the tissue-level, leaving the question unanswered as to which cellular or sub-cellular ‘regulatory level’ selection may act upon. In 2017, we applied single-mt sequencing to primary cultures of mouse and human neurons and astrocytes and found allelic variation within a single mitochondrion over the multiple genomes encapsulated in a single organelle. We showed larger inter- and intra-cellular single mt allele frequency (AF) variation in mouse than in human neurons, suggesting that the two species may have evolved different ways for mt segregation (22). A limitation of this work was the small sample size for individual mitochondrion due to manual organelle collection, which limited the ability to estimate the intra-cellular mt variation.

To generate the single mt landscape of single-nucleotide variants (SNVs) in specific regions from dispersed cell cultures of mouse cortical brain region astrocytes and neurons, we developed a highly scalable moderate throughput single cell -single mitochondria isolation procedure using fluorescence-activated mt sorting of mitochondria-associated TOMM20 antibody conjugated microbeads (23, 24). Our approach has allowed us to study inter- as well as intra- cellular mtDNA variation from isolating an average of 10 times more single mitochondria from single neurons and astrocytes than previously reported (22). Briefly, mouse neurons and astrocytes from brain cortex dissection were seeded on coverslips and maintained in primary neuronal cell culture at 37°C, 5% CO_2_. The coverslips were stained with MitoTracker Red before the single neurons and astrocytes were harvested using micropipette and transferred into an Eppendorf tube. The single cells lysates were incubated with anti-TOMM20 conjugated microbeads for flow cytometry of captured mitochondria from single cells (see methods for details and Figure 1A and Figure S1A for number of mitochondria isolated per cell and per mouse pup). After sorting single mitochondria into 96 well plate, we developed a two-prong approach to amplify mt-genomes from individual single isolated mitochondria. We adapted Rolling Circle Amplification (RCA, linear amplification method) to amplify individual mt-genomes from single mitochondria producing yields large enough to perform multiple subsequent PCRs. The mitochondria ID barcoded PCR primers were designed to amplify 12 target regions containing previously reported SNVs (22) (Table 1). Using this method, we were able to detect a spectrum of known and novel SNVs from over 1,645 single mitochondria from ∼100 single astrocytes and neurons isolated from 13 mice in our single-mitochondria (SMITO) dataset. We identified 1032 SNVs showing prevalence bias based on the target regions to which they belong. Both somatic and inherited mt variants are associated with diseases. Hence, these were further classified into inherited or somatic SNVs to elucidate the influence of evolutionary processes on heteroplasmy dynamics. Comparison of these two classes of SNVs, showed distinct regional differences in the occurrence and transition-transversion bias. For the inherited SNVs, AF variation at the levels of single mitochondria, cell, and animal revealed overall variation to be the largest at the animal level. Also, specific SNV sites exhibited relatively higher inter- and intra-cellular variation as compared to the variation at the animal level. Apart from the global role of mitochondria in a cell, it is known that mitochondrial metabolism in neurons vs astrocytes is different having evolved to meet specialized roles of neurons and astrocytes (25, 26). We found that astrocytes exhibit more SNVs than neurons and two SNVs with differential incidence, suggesting distinct mutational propensity in these two cell types and purifying selection in neurons. In addition, we detected linked SNV pairs within the analyzed regions that suggests co-segregation.

**Figure 1.**
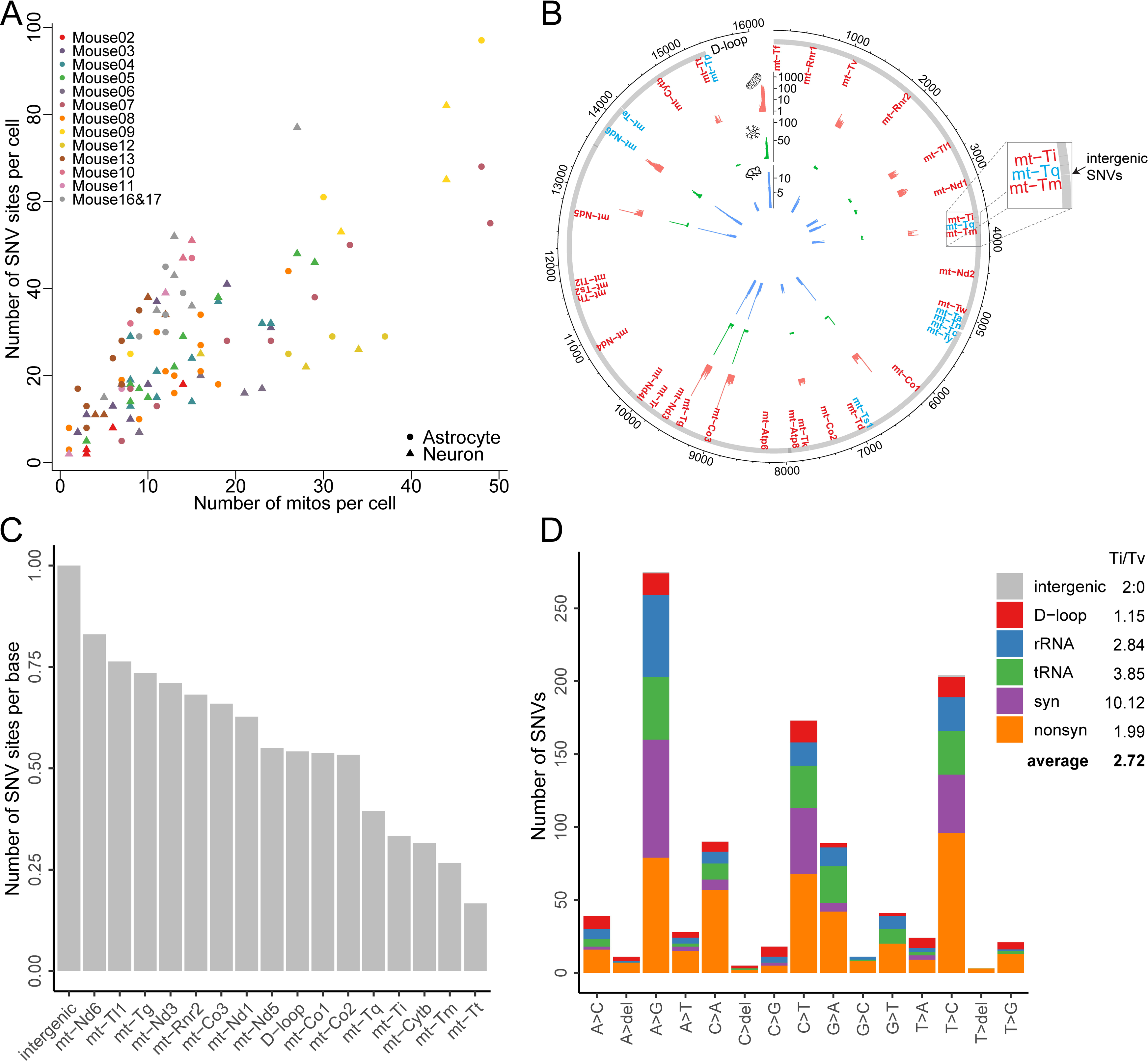
SNV distribution in SMITO dataset. (A) The number of SNVs sites per cell as a function of the number of mitochondria in each cell. Mitochondria collected from each mouse are color coded, from astrocytes and neurons are depicted by circles and triangles respectively. (B) Circos plot showing distribution and site of detected SNVs along the mitochondrial (mt)-genome. The bar height indicates the number of mitochondria (red), cells (green) and mice (blue). Thirty-seven mt genes are marked along the thick circumference (red denotes H-strand genes, blue indicates L-strand genes). (C) The number of SNVs per gene/region normalized to SMITO- analyzed target region length (bp). (D) A SNV mutational spectrum with the number of SNVs on the y-axis grouped by base substitutions: transitions (Ti) and transversions (Tv) on the x-axis. Ti and Tv are color-coded by region (D-loop, rRNA, tRNA) or functional impact in protein-coding region (synonymous, nonsynonymous).

**Table 1.**
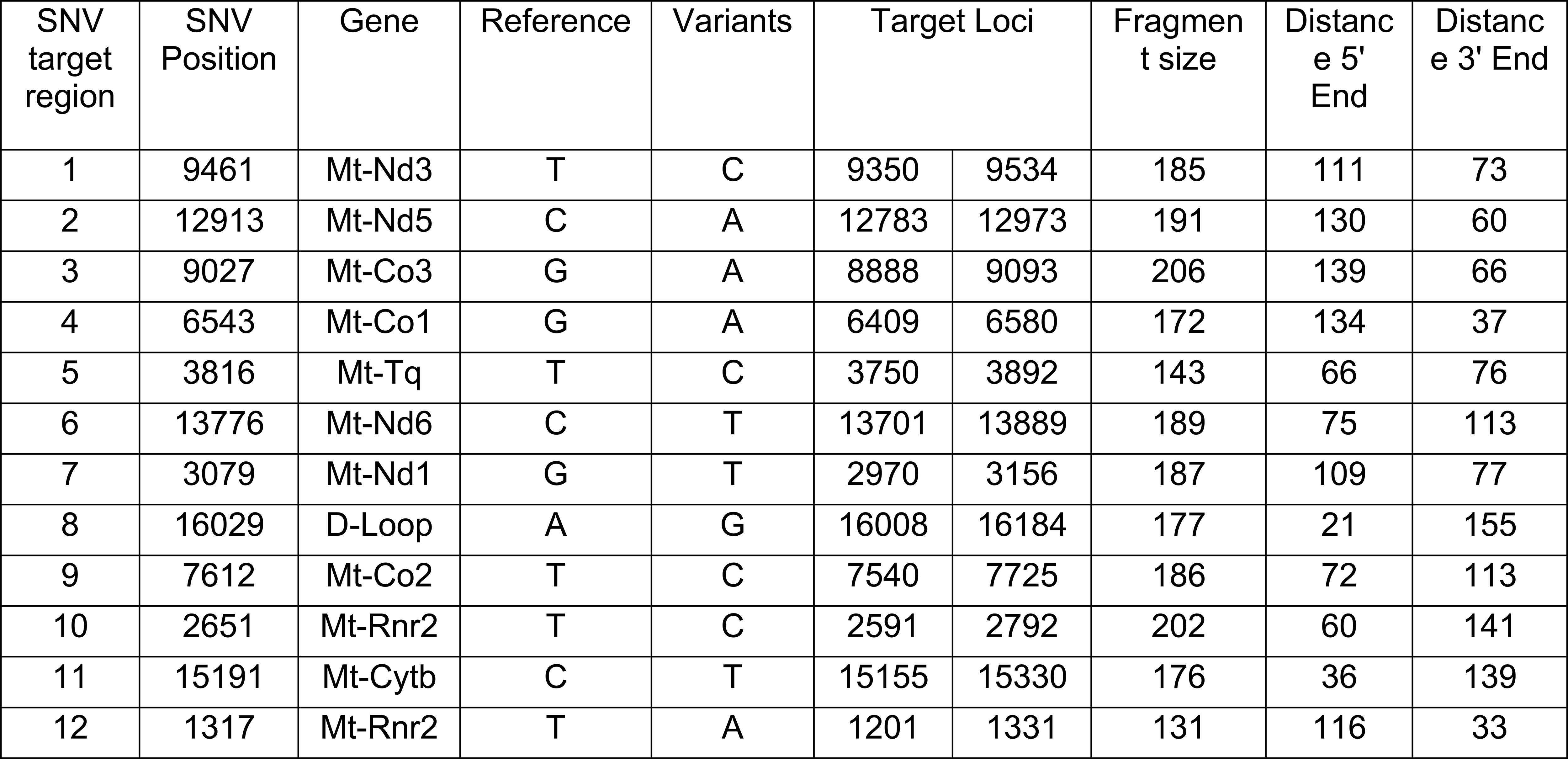
List of SNV target regions.

## Results

### SMITO SNV detection, distribution, and quality assessment in each mitochondrion sample

After QC filtering of data collected from 1645 mitochondria, 102 cells and 13 mice (the co-cultured Mouse16&17 regarded as one animal; the pup brains in such cases are trypsinized together and then plated to give rise to these mixed pup cultures that were then maintained in culture for days) (Figure S1, Dataset S1), 1032 SNVs distributed over 838 unique sites were obtained across 12 target regions (Table 1 lists the target regions as well as a previously reported SNV within each region); 1519 mitochondria showed at least one SNV. Out of the 12 SNVs specifically targeted from previously reported SNVs (22), we found 4 in our new samples: 9027:G>A (in 1308 mitochondria, 100 cells, 13 mice), 9461:T>C (in 1198 mitochondria, 100 cells, 13 mice), 15191:C>T (in two mitochondria, two cells, two mice), and 3816:T>C (in only one mitochondrion). We detected 4 of the 12 SNVs that were reported in Morris *et al*. (2017) potentially due to differences in mouse population as well as neuronal cell populations (hippocampal cells were also used in the previous study as opposed to only cortical neuronal cultures used for the SMITO study. Among previously reported SNVs (22) (*n* = 285), 35 are within the sequencing interval of SMITO and 22/35 (63%) SNVs were observed by our SMITO data. On average, we detected 4.5 variant sites within the targeted regions (∼1/10^th^ of genome) per mitochondrion (Figure S2) and the number of variant sites per cell scales linearly with the mitochondria count in a single cell (Figure 1A). The SNV count was found to be significantly associated with the mouse identity (ANOVA *p* < 2e-16), suggesting that mice can have different amounts of mutation load, likely through differential inheritance and total somatic mutations.

Figure 1B shows the distribution of SNVs with respect to mt-genes and the number of mitochondria, cells, and mice sharing each variant site. After our filtering criteria (see Methods), most (75%) of the SNV positions were found to be shared by at most three mitochondria (three cells or two mice). The Region 8 in the D-loop (16008-16184) showed most abundant SNVs. The observed spikes at SNVs 6430, 6432, 9027, 9461 and 12831, indicates highly ubiquitous SNVs. Single-mt-SNVs at 6430 and 6432 were prevalent in a subset of 5 mice. The width of the shaded patches in Figure 1B indicate dispersion of the SNVs within a specific target region like in region 1 (9350-9534, mt-ND3), 3 (8888-9093, mt-Co3), 6 (13701-13889, mt-Nd6) and 7 (2970-3156, mt-Nd1). The intergenic target region contained the highest number of SNV sites per base (=1 as a result of 2 SNVs for 2 bases coverage), followed by the region in mt-Nd6 (98 sites for 118 bases) as shown in Figure 1C.

We further analyzed the spectrum of mt mutations. Among the 1032 unique SNVs in total (Figure 1D), transitions (A<>G, T<>C) predominated base substitutions (Figure S3), consistent with findings in mouse brain and muscle (16). Our analysis reveals that the occurrences of C>T and G>A changes were as frequent as T>C and A>G changes (Figure S3) indicating that deamination errors (27) were unlikely to affect our dataset. There are 440 nonsynonymous, 189 synonymous, 161 tRNA, 147 rRNA and 93 D-loop as well as two intergenic (between mt-Tq and mt-Tm) SNVs (Figure 1D). Among these, 42 are predicted to be high impact (generating stop codon or reading frame shifted) (Table 2) and Dataset S2 lists the potential functional impact of these 1032 SNVs along with SIFT (Sorting Intolerant from Tolerant) (28–30) prediction (whether an amino acid replacement has an impact on protein functionality) results. Using the SIFT prediction for non-synonymous SNVs, we tested if greater number of mitochondria and cells shared “tolerated” SNVs as compared to the “deleterious” SNVs using Wilcoxon’s rank-sum test (Figure S4). However, the effect size was minor and no statistical significance was detected (including Poisson regression test).

**Table 2.**
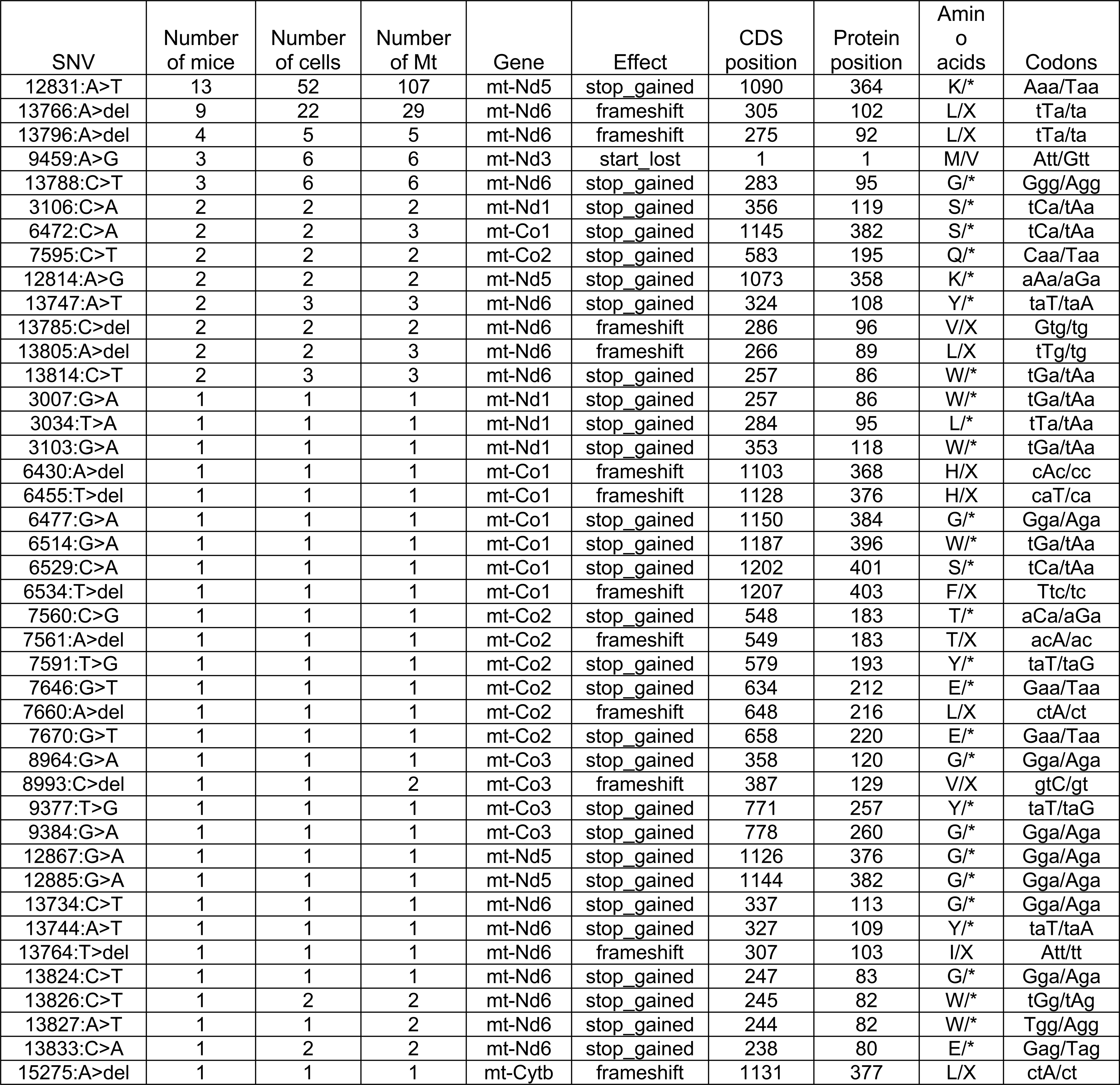
List of the high-impact SNVs annotated by Variant Effect Predictor.

We also checked SNV impact on tRNAs, focusing on hits in the anticodon region. In total, there were 41 SNVs (at 40 unique sites) affecting tRNAs; among them, 24 hit mt-Tg, 12 hit mt-Tl1, 3 hit mt-Tm and 2 hit mt-Tq. By assessing distance from SNV to anticodon center, a single candidate, 9423:C>T was identified, which would impact the 3rd position of the anticodon TCC> TCT (Figure S5). The mutated tRNA may still be able to recognize the codon GGA because U can form a wobble pair with G. Since the vertebrate mitochondrial stop codon is AGR (R=A/G), we reasoned that 9423:C>T might rescue premature termination caused by nonsense mutations (e.g. 6472:C>A, 12831:A>T). Yet, the low frequency of 9423:C>T (observed in only 4 mitochondria from 4 cells in 3 mice) hindered further association testing.

### Mouse strain associated segregating sites and inherited-somatic SNV classification

Since each mitochondrion has multiple genomes, the frequency of each base-pair variant was estimated by read frequency and adapting the population genetics terminology, called “allele frequency” (AF). The variant AF (VAF) distribution for all SNV sites showed a U-shape with the two modes in the range of 5-10% and 95-100% (Figure 2A), reflecting potentially neutral drift of majority of the allelic variants with 0 and 1 absorbing states. We also compared SNV sites (838) from our SMITO dataset against the segregating sites from the mitochondria of 17 mouse strains and observed a significant (hypergeometric test *p* = 0.0034) overlap (*n* = 89) (Figure 2B). Assuming the 17 strains were an unbiased sample, the AF averaged over 17 strains was calculated and the highest AF was selected as the major allele (Dataset S3) and the second highest AF as the top minor allele. Among the 89 overlapping sites, we observed a 98% match for major alleles, consistent with segregating variation seen in the strains and a 63% match for minor alleles. In theory, each site can have up to 3 minor alleles. For example, a major allele ‘A’ can have C,G, T minor alleles or a deletion and the expected match should be merely 25% (1/4) by chance.

**Figure 2.**
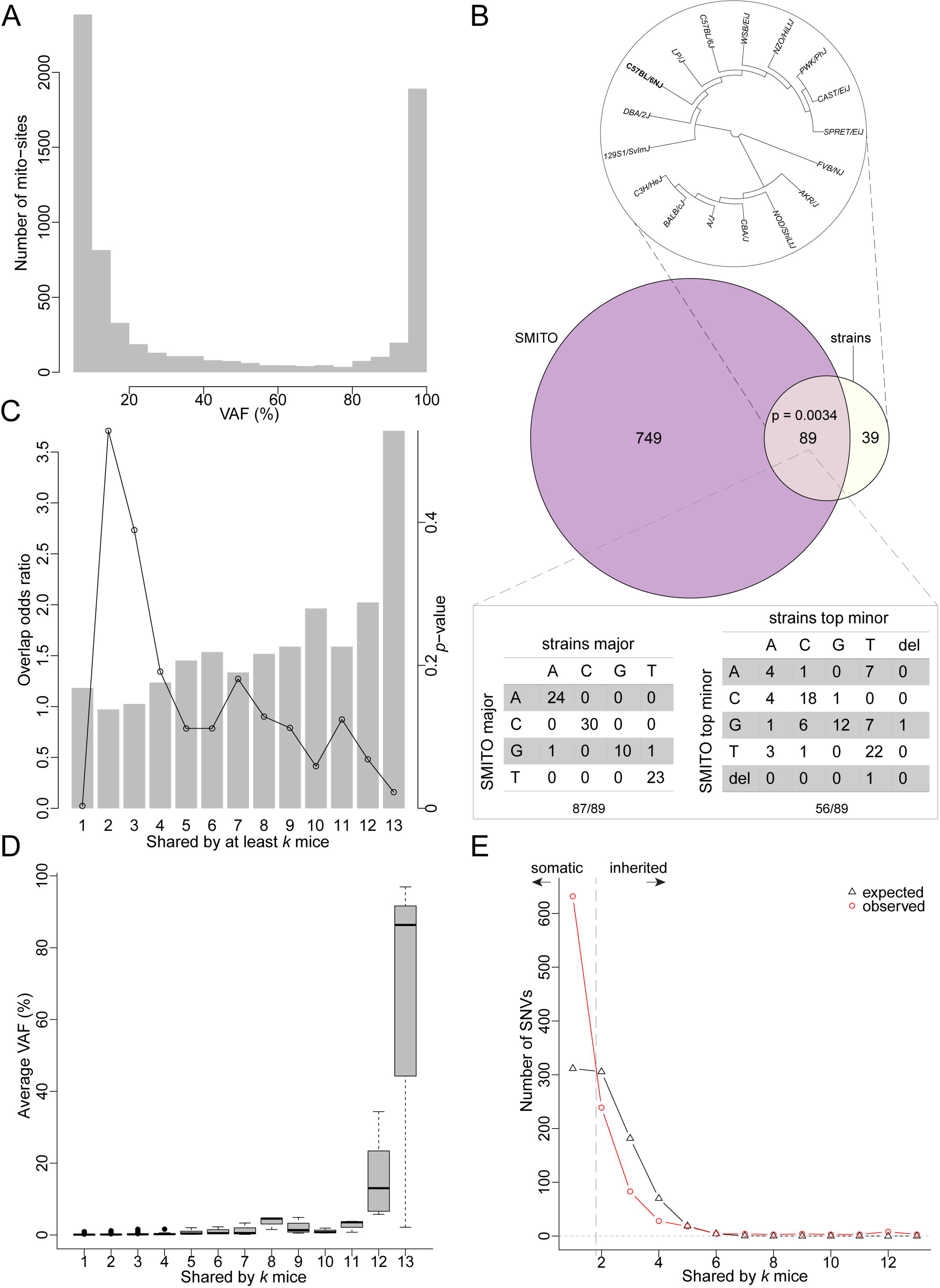
Mouse strain associated segregating sites and inherited-somatic SNV classification. (A) Distribution of the variant allele frequencies (VAFs) on the x-axis for all the variants/SNVs observed in all the single mitochondria samples on the y-axis. (B) Top: the neighbor joining tree built on 17 mouse strains’ segregating sites; middle: the Venn diagram showing the overlap between SMITO and the mouse strains data limited to the 2145 bp analyzed in this study; bottom: consistency in the major alleles (left) and top minor alleles (right) between SMITO and the mouse strains data. (C) The barplot shows the strength of the overlap of the mouse strains and SMITO data conditioned on the number of mice shared. The dotplot shows the *p*-value from hypergeometric tests. (D) The relationship between average AFs on the y-axis and the number of mice shared on the x-axis. (E) Classification of somatic and inherited SNVs by the theoretical (black triangles) and the observed (red circles) distribution of the SNV count (y-axis) as a function of the number of mice shared (x-axis).

Given the lower likelihood of somatic mutations being shared between animals, we hypothesized that a higher prevalence of an SNV shared amongst animals increases the likelihood of it being inherited. Of our 838 unique SMITO sites, the non-overlapping 749 sites were most likely not derived from germline inheritance. To assess the overlap of SNVs from the mouse strain data with our SMITO dataset, we calculated the overlap odds ratio as a function of the number of mice sharing SNVs. Using the mouse strains data as a plausible pool of inherited SNVs, we observed stronger overlap when thresholding the SNVs by higher number of mice that share the SNV (Figure 2C) as seen by the bars that depict the odds ratio and the *p*-value depicted by the dotplot within the same graph. According to the “drifting model” of neutral alleles, the “older” an SNV, the more extreme AF (i.e., towards 0 or 1). As anticipated, we observed increasingly larger AF variation when the number of mice shared went higher (Figure 2D). We also noticed a monotonic increase in the average AF especially as the number of mice sharing SNVs exceeded 9. This trend was also true for the number of cells or mitochondria sharing SNVs (Figure S6A-B). The three SNVs found to be shared by all 13 mice were 9027:G>A (average AF 86.3%), 9461:T>C (average AF 96.9%) and 12831:A>T (average AF 2.2%). Less shared SNVs are presumably “young” and are expected to have lower frequency as compared to more highly shared SNVs.

Using the hypothesis that shared SNVs among mice likely indicate inherited rather than de novo mutations, we categorized SNVs as somatic or inherited. The number of SNVs as a function of the number of mice that shared the variant was modelled (Figure 2E, S3). A null distribution was generated where SNVs were assumed random while SNVs per mouse were maintained. We found several SNVs unique to an individual mouse but a lower-than-expected SNVs shared by 2-4 mice (Figure 2E). From this analysis, the SNVs unique to each mouse were designated as “somatic”, and the SNVs shared by ≥ 3 mice as “inherited” for the rigorous analysis. As a result, we classified 632 SNVs as somatic and 161 as inherited SNVs.

### Mutational preference comparison between inherited and somatic SNVs

We next compared inherited and somatic SNVs for their mutational pattern. The two classes showed distinct regional preferences for transitions (Ti, purine to purine or pyrimidine to pyrimidine change) and transversions (Tv, purine to pyrimidine change or vice versa). Inherited SNVs occurred most frequently in D-loop (20%) and mt-Nd6 (19%), while somatic SNVs occurred most frequently in mt-Rnr2 (15%); only 8% and 11% somatic SNVs were found in D-loop and mt-Nd6, respectively (Figure 3A). Somatic SNVs within mt-Co1, mt-Co2, mt-Tq, and mt-Tm were detected two-times more often than inherited SNVs. To control for the PCR coverage for each region, we normalized the SNV occurrence per mitochondrion by the actual number of bases targeted by SMITO within each gene and defined it as the per-base variant rate. The somatic variant rate ranged from 0.16 to 0.51 while the inherited variant rate ranges from 0.02 to 0.23 (Figure S7), derived by the absolute number of SNVs (∼4 times more somatic than inherited SNVs). We observed a stark difference in the variant rate distribution (Figure 3B). Inherited SNVs exhibited an uneven distribution when compared to the average, variants were more prevalent in mt-Tg, D-loop, mt-Nd6, mt-Nd3 and mt-Tl1 but less prevalent in mt-Tq, mt-Tm, mt-Co2, mt-Cytb, and mt-Nd5. In contrast, somatic SNVs showed a more uniform distribution as expected; only a modest decrease was seen in mt-Tt (Figure 3B). The fact multiple animals have the same SNV, despite germline bottleneck, suggests some kind of maintenance mechanism.

**Figure 3.**
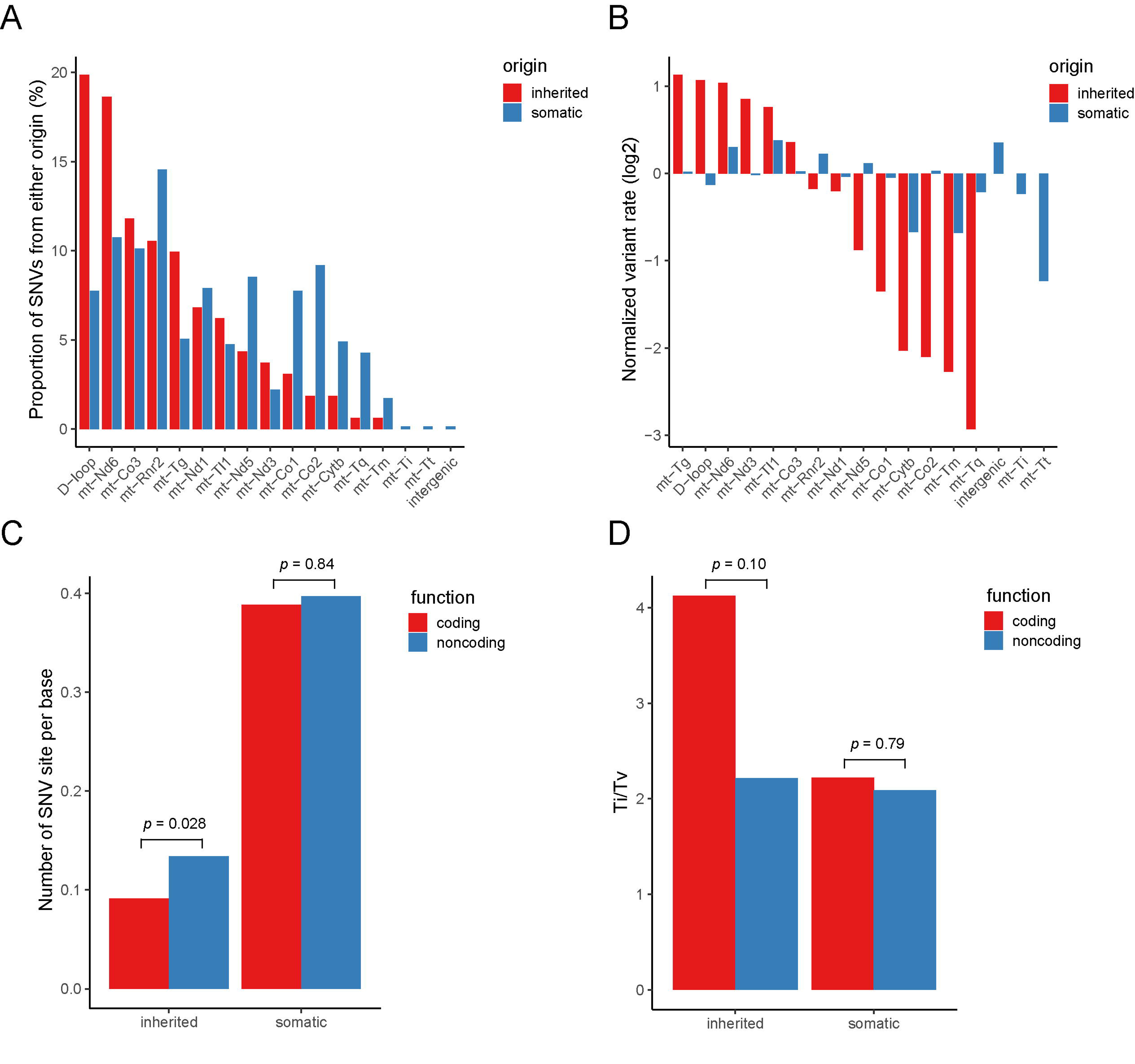
Mutational preference comparison between the inherited and somatic SNVs. (A) The proportion of inherited (red) and somatic (blue) SNVs within target region (gene loci). The percentages were calculated within each SNV class: 161 for inherited and 632 for somatic. (B) Length-normalized mean-centered variant rate for inherited (red) and somatic (blue) SNVs in each gene/locus. (C) The number of SNV sites per base in coding (red) and non-coding (blue) regions for inherited and somatic group of SNVs. (D) The Ti/Tv ratio of inherited and somatic SNVs classified by coding (red) and non-coding (blue) regions.

We compared the per-base variant rate for inherited and somatic SNVs in both coding and non-coding regions (Figure 3C). A marginal significance (p=0.028) was observed between coding and non-coding regions for inherited group SNVs but not for somatic ones, confirming the observation in Figure 3B that somatic SNVs had more uniform, functionally irrespective distribution. Similar comparison between coding and non-coding regions was made for Ti/Tv (Figure 3D). Although neither showed p<0.05, Fisher’s combined p=0.015 hinted difference between the two classes of SNVs.

Ti/Tv ratios were compared for two classes of SNVs, revealing significant differences based on target regions and functional categories (Figure S7C-D). Inherited SNVs displayed higher Ti/Tv ratios in mt-Co2, mt-Cytb, mt-Nd3, mt-Rnr2, mt-Tl1, and mt-Tm, indicating a prevalence of transitions. Conversely, somatic SNVs generally exhibited Ti/Tv ratios < 2.5 in these regions. However, mt-Tq, mt-Nd1, mt-Tg, mt-Nd5, and mt-Co3 showed inherited SNVs with Ti/Tv ratios twice as high as somatic SNVs. Notably, mt-Co1 was the sole protein coding gene displaying a fourfold higher Ti/Tv ratio in somatic compared to inherited SNVs. Fisher’s exact tests revealed that inherited SNVs had higher Ti/Tv ratios in rRNA and tRNA (p = 0.001 and 0.02, respectively), while somatic SNVs had higher Ti/Tv ratios in the D-loop (p=0.002) (Figure S7D). On comparing the Ti/Tv values from the polymorphisms from between -genera, -species (Figure S8A), -strains (Mouse Genomes Project) and -population data (Figure S8B), we observed a clear trend for the whole mt-DNA (Figure S9A): (1) phylogenetically more distant mt-genomes showed lower Ti/Tv, (2) D-loop had the lowest Ti/Tv while synonymous polymorphisms had the highest Ti/Tv. When limiting the polymorphisms to the SMITO assayed regions (Figure S9B), we observed again low Ti/Tv values in nonsynonymous and D-loop polymorphisms but elevated Ti/Tv in synonymous polymorphisms in between-genera, -species, -population (Dataset S4). We used the McDonald-Kreitman test on mouse population data to test if evolutionarily selected genes show differential Ti/Tv from neutral genes. However, no gene showed significance for selection (Dataset S5).

### Hierarchical distribution of the AF variation and heteroplasmy of inherited SNVs

New mutations in the mt-genome alter frequency through replication, drift, and selection. AF changes may occur due to unequal nucleoid replication within a single mitochondrion, uneven genome segregation between mitochondria (fission/fusion), segregation of mitochondria during cell division at the cell level, and germline transmission of inherited SNVs at the animal level. Thus, AF can be modulated at multiple levels. To investigate the variation of AF from a hierarchical perspective, we estimated the AF variation for inherited SNVs at three levels: between -mice, -cells, and -mitochondria using a nested ANOVA model after arcsine transformation (for details see Methods). To make variations across levels comparable, the mean-of-squares (MS) were measured. Most of the SNVs showed larger (average of 3 times) variations at the mouse level than the cell level (Figure 4A, left), and a larger (average of 1.4 times) variation at the cell level than the mt level (Figure 4A, right). These systematic biases are unlikely to be experimental or biological noise (Figure S10) since random noise is expected to result in an equal MS across all three levels. The mouse and the cell identities were shuffled to generate a null distribution of the MS at the three levels. The null distribution level showed an average of 1/3, while the mouse level showed twice the amount of AF variation (Figure S10A). SNVs according to the between-mice variation were plotted in a descending order (Figure 4B). SNVs with the top between-mice AF variation include 6430:G>C, 6432:A>C, and 9027:G>A (Figure S10B) whereas SNVs 2659: C>T, 15266: C>T and 1253: A>G SNVs exhibit larger inter- and intra-cellular AF variance. These data suggests that heteroplasmy observed for different sites may be a result of site-specific selection.

**Figure 4.**
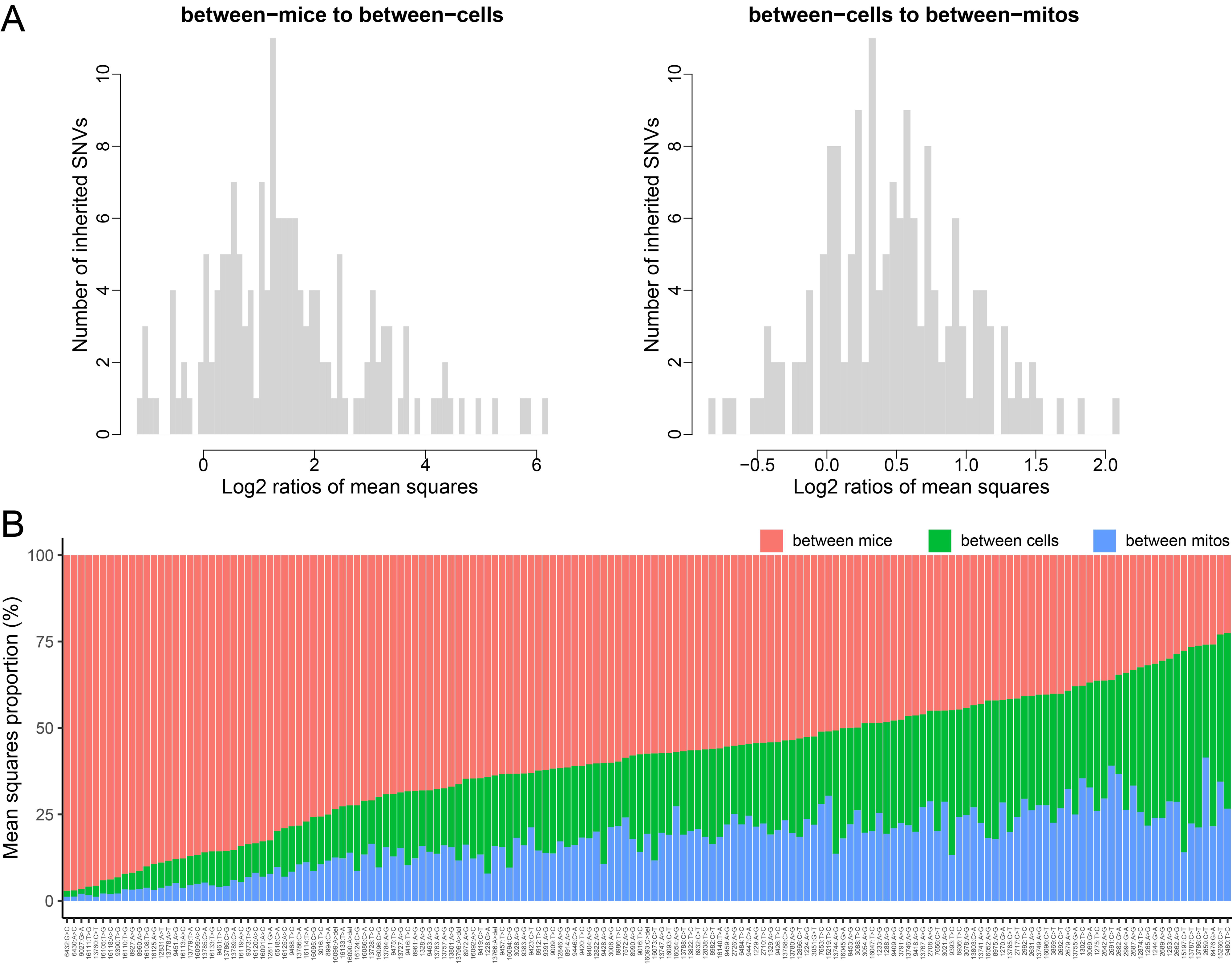
Comparison of the inherited SNVs’ allele frequency (AF) variation across mouse, cell, and mitochondria level. (A) The distribution of the ratios of: between-mice to between-cells variation (left) and between-cells to between-mitochondria variation (right). (B) The proportion of between-mice (red), between-cells (green), and between-mitochondria variation (blue) for each inherited SNV depicted on the x-axis.

### Comparison between astrocytes and neurons in the SNV incidence and AF variance

Unlike neurons, astrocytes are mitotic and are known to have more reactive oxygen species due to lower mitochondrial bioenergetic efficiency (25). We hypothesized that astrocytes would exhibit more SNVs than neurons. On comparing the number of SNVs presented in the two cell types, we observed a mild yet significant difference (*p* = 0.024, for details see Methods) between the two cell types, with neurons showing ∼5% lesser number of total SNVs when compared with astrocytes (Figure 5A). We further investigated if there was cell-type difference in the SNV occurrence for specific SNV(s). By fitting the presence or absence of an SNV to a logistic regression model (the mouse effect included as the covariate), we found that 9419: C>T occurred more frequently in astrocytes and 9027: G>A more frequently to neurons with significances of *p* = 0.035 and 0.038, respectively (Dataset S6). As an independent test, we compared the fraction of SNV-carrying mitochondria between the two cell types controlling for the mouse effect and we were able to recapitulate significance of these two SNV differences (*p* = 0.036 and 0.052, respectively) (Figure 5B). To understand the functional consequences of 9419:C>T which impacts the first base of the Ac-loop of mt-Tg (31), we used RNAfold (32, 33). RNAfold predicted this mutation to cause the minimal free energy to decrease from -6.60 kcal/mol to -8.60 kcal/mol, which may result from an additional A:U pair preceding the anticodon (Figure S11). We found that 9027:G>A showed twice as much between-mt AF variance in astrocytes than in neurons (Figure 5C). The cell specific differences in the percentage of mitochondria with certain SNV site alleles (9419: C>T and 9027: G>A) emphasize differences in maintenance of SNVs for mt- genome between neurons and astrocytes. To further investigate if evolutionary constraints on mt protein coding genes were different between neurons and astrocytes, we performed Ka/Ks analysis which takes into account the ratio of the number of nonsynonymous substitutions per nonsynonymous site (Ka) to the number of synonymous substitutions per synonymous site (Ks). Ka/Ks =1 suggests neutrality, <1 suggests purifying selection and >1 suggests positive selection. We also found a reduced Ka/Ks relative to the background distribution (*p*=0.04), indicating an overall purifying selection (Figure S12 and Dataset S7). Interestingly, astrocytes did not show significance for purifying selection whereas neurons did (*p*=0.01). A recent study by Hu *et al.* (34) shows that neuronal cells are under strongest evolutionary constraint and higher selective pressure as compared to the other somatic cells including astrocytes, by single-cell RNA seq data analysis. Since neurons are the functional units of the nervous system and have limited regeneration, have a high energy demand, and are post-mitotic, the selection pressure experienced by these cells is higher than compared to mitotic cell types like astrocytes. (Figure 5D). But we also note that since neurons are post mitotic cells, the effective population size of non-dividing cells might be smaller than dividing cells and hence genetic drift in neurons may influence the changes in allele frequencies.

**Figure 5.**
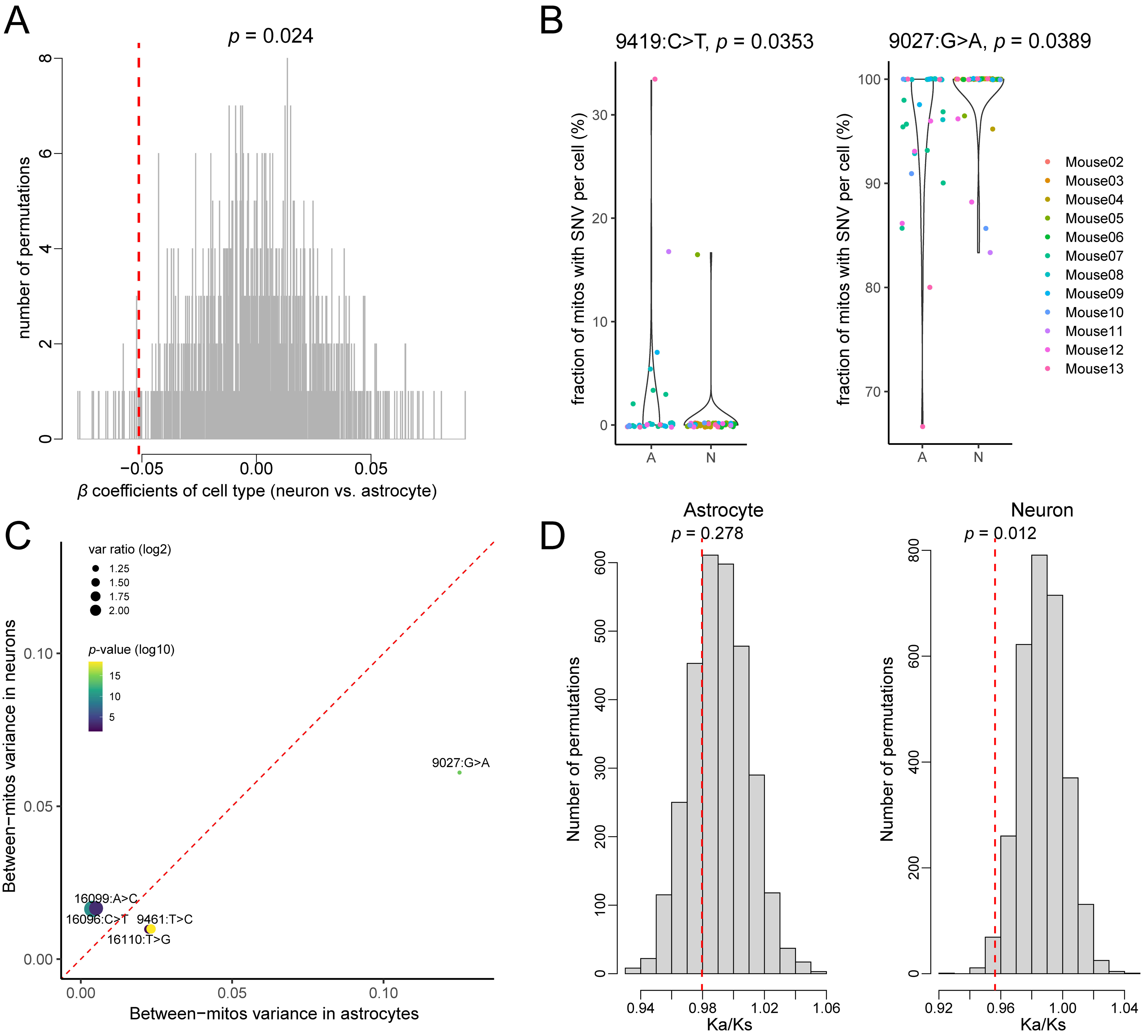
Cell type difference in the SNV incidence and allele frequency (AF) variance. (A) The null distribution (gray lines) and the observed value (red dashed line) of the cell-type effects on the number of total SNVs per mitochondrion. (B). The percentage of SNV-carrying mitochondria per cell for astrocytes ‘A’ or neurons ‘N’ for 9419:C>T (left) and 9027:G>A (right). (C) The scatterplot depicting the between-mitochondria variance in neurons on the y-axis to the between-mitochondria variance in astrocytes on the x-axis. The size of circles indicates the ratio of the variance (absolute value of log2), and the color gradient indicates the *p*-value (minus log10) from F-test (two-sided). (D) Ka/Ks statistics in astrocyte SNVs, and neuron SNVs. The red dashed line represents the observed Ka/Ks in respective cell type.

### Linkage of SNVs in the mt-genome

We identified 9 pairs of SNVs within the target regions showing significant (BH adjusted *p* < 0.1) linkage across all cells and mice at position 6430, 6432, 9027, 9461, 16108, 16111, 16118, 16133 and 16140 (Figure 6A), most of which were found within the D-loop region. Interestingly, two high AF SNV sites 9027 and 9461 showed strong linkage (Fisher’s exact test *p* = 8.82e-5). Linkages can result from (1) co-segregated mitochondria or (2) an absence of recombination. Some short-range linkages such as 6430-6432 shared at single-cell level, in 3 cells from mouse 12 and 16-17, were caused by the lack of recombination, as the variant alleles were located at the same read (haplotype) (Figure 6B, Figure S13).

**Figure 6.**
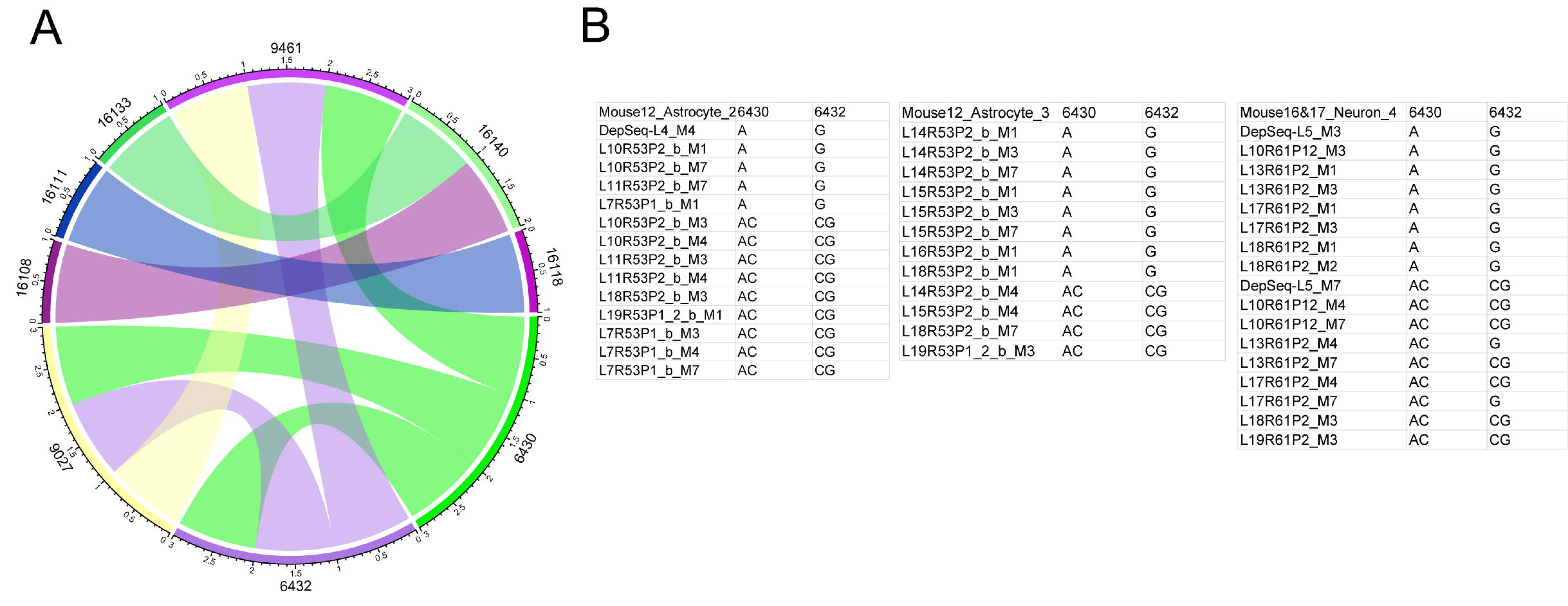
Linkage analysis of mt-SNVs. (A) Circle chord graph depicting significantly linked SNV loci, BH corrected *p-*value < 0.1. (B) Single-cell level significant linkages with genotype shown in each mitochondrion from each of 3 individual cells.

## Discussion

This study addresses the mt-SNV variance at the mouse, cellular and single-mt levels, heteroplasmy of specific SNV loci, cell type differences, and SNV segregation in the mouse neurons and astrocytes. There are differences in the prevalence or distribution of variants from specific regions of mt-genome because of their potential impact on the mitochondria or the cell. Given that we only targeted ∼1/10^th^ of the mt-genome, the number of SNV sites per mitochondrion found were relatively high compared to our previous study. This may be a result of larger sample size (∼13 fold) at the level of individual mitochondria. This may also indicate that besides D-loop, there are certain regions in the mt-genome like rRNA/tRNA (as compared to the other target regions analyzed) are more permissive to a nucleotide change (Fig S1A). Previously, single-cell mtDNA mutation analysis in lymphocytes and monocytes from a 76 year-old female revealed that 70% of the mtDNA mutations were non-synonymous (35). Similarly, we found that the 12 target regions analyzed had greater number of non-synonymous SNVs (∼70% of these were transitions) as compared to the synonymous ones. Our comparison with mt sequences from related mouse strains and species suggests that a large fraction of our SNV sites (evolutionarily shared 89 sites) might have been ancestrally inherited, while the rest of SNVs might be *de novo* mutations whose effects might be recessive to other nucleoids in the same organelle, which explains the overall high proportion of the nonsynonymous changes in the target regions we analyzed.

Three SNVs were shared by all 13 mice: namely, 9027:G>A (average AF 86.3%), 9461:T>C (average AF 96.9%), and 12831:A>T (average AF 2.2%). Site 9461: T>C elicits a change in the start codon (AUU to AUC) for mt-Nd3 (36). 9027:G>A causes a missense (glycine-to-serine) change in mt-Co3 and 12831:A>T results in premature truncation of mt-Nd5 (after∼60% of mRNA has been translated). These three sites show SNVs common to all sampled animals, despite germline bottlenecks in each generation, suggesting that they are actively maintained amongst mice. At 9461 position, two strains (PWK/PhJ and C57BL/6J) use T while 15 others use C. We previously found that this site is also polymorphic amongst mouse strains (22), which was hypothesized to be cryptic heteroplasmy obscured by bulk sequencing. Thus, we continue to reason that 9461 is ancestrally polymorphic and under balancing selection in macro-evolutionary scale.

The overall nonsynonymous-to-synonymous ratio is approximately 2, suggesting a large fraction of the observed SNVs are randomly distributed over codon positions and transiting through purifying selection or are selectively neutral. Synonymous SNVs showed the highest transition-to-transversion (Ti/Tv) ratio (∼10), which may occur as a result of wobble position degeneracy and/or RNA secondary structure constraints (36); tRNA and rRNA showed high Ti/Tv (3.8 and 2.8, respectively), while D-loop showed the lowest Ti/Tv (∼1). The difference of Ti/Tv between tRNA, rRNA and D-loop possibly indicates different tolerance to changes in the thermal energy of base pairing due to the requirement of rRNA structure for translation while the D-loop region is gene regulatory and may be permissive to transversions that may promote regulatory protein binding to DNA. The observed transition bias (Ti/Tv of 2.72) in the mouse mtDNA is in agreement with previous studies (37–39). Apart from the protein coding regions analyzed, the higher transition bias for the rRNA and tRNA target regions suggests that purification selection is not limited to protein coding regions in mouse mtDNA. However, since the transition transversion ratio varies between species, it will be interesting to see if Ti/Tv ratio for human mtDNA have significant differences between regulatory regions versus protein coding regions.

We found that the variants in the coding regions classified as ‘inherited’ were mostly biased towards transitions over transversions. Interestingly, we found that that after the D-loop, which is known to be hypervariable, the number of transversions was highest in the mt-Nd6, a large fraction of which were non-synonymous. We found that SNVs in ND6 were predicted to have a higher impact when compared to other regions analyzed as noted in Table 2. Intriguingly, a study in 2013 by Bannwarth *et al*. reported ND6 and ND5 were the two protein coding genes in the mt-genome that had the most mutations in patients with mitochondrial diseases (40). Being a crucial component for oxidative phosphorylation (OXPHOS), mt-Nd6 SNVs exhibiting a deleterious impact may result in a need for increased replication and transcription as a compensatory response, as observed in mt complex I from human skeletal muscle (41). Thus, deleterious sequence variants within a gene may lead to clonal expansion of ‘imperfect’ mt-Nd6 or complex I genes until a threshold is reached, as a result of compensatory increase in mtDNA replication (42, 43).

The hierarchical extent of overall variation shows the greatest variation between animals, followed by between cells, and the least variation manifest between mitochondria within a cell. This most likely reflects mainly neutral drift following the overall demographic structure, where the mt lineages have the highest separation between animals, then cells, then organelles within the same cell. At the level of cells, additional homogenizing forces can reduce variation such as suggested by Twig *et al.* that in mouse islet cells mitochondrial fusion and fission are coupled events (44). Our data also suggests that there are some SNVs that show negligible variance at the mouse level but exhibit large variance between mitochondria within a cell i.e., heteroplasmy. Like the codon position-specific selection pressure observed in mtDNA germline variants (45), this points to a region/gene loci-specific mechanism of SNV maintenance in a cell lineage for particular regions of mt-genome. The differences in mt-SNVs variance at cellular vs mt level at specific sites, is important to consider for therapeutic strategies. In diseases with increased cellular variance of pathogenic SNVs, targeting and eliminating cells with higher mt-SNV prevalence could be a potential treatment.

The mitochondrial metabolic pathways in neurons and astrocytes are prioritized by their respective key function in the central nervous system, i.e. neurotransmission by neurons and support homeostasis in neurons by astrocytes. Several studies have shown that a main mt difference is that neurons are primarily dependent on OXPHOS to meet their high energy required for neurotransmission, however, astrocytes are primarily dependent on glycolysis. As a result of this main distinction, there are consequential differences in the bioenergetics, maintaining ROS, and signaling. An example of this is prioritizing conversion of pyruvate to lactate in order to shuttle lactate to neurons (since neurons are extremely efficient in utilizing lactate for OXPHOS) (25, 26, 46). Since these two cell types are heavily interdependent, it is understandable that dysfunction in one cell type will impact the functioning of the other and has implication in numerous neurodegenerative disorders (47–49). In this study, we have showed that there are cell type specific differences in the ways mt-SNVs accumulate and propagate in these cells. The greater number of SNV sites in astrocytes (mitotic) than neurons (nonmitotic) is in agreement with the concept that mitotic cells tend to acquire more somatic mt mutations (42) and that astrocytes tend to have higher ROS production as compared to neurons (46, 50). Given the subpopulation of mitochondria and specific target regions analyzed in the present study, it is likely that this difference will be more pronounced when the entire mt-genome is screened. Mt-SNV 9027: G>A is prevalent in multiple mouse strains, but no cell specific differences have been previously reported. This study demonstrates for the first time that even in a wildtype mouse pup, there exist differences in the variance of allele frequencies and levels of heteroplasmy of certain SNV sites between neurons and astrocytes. For both sites 9419: C>T and 9027: G>A, the mitochondria analyzed in a neuron showed less variability as compared to astrocytes. This may be explained by segregation during mitosis in astrocytes which is absent for post-mitotic cells including neurons as well as stronger purifying selection noted in neurons (Fig 5D). The purifying selection of mt- SNVs in neurons as shown by the Ka/Ks statistic suggest that neurons are under stronger evolutionary constraint as compared to astrocytes. Functionally, 9419: C>T is a mutation in tRNA glycine which leads to a RNA structural change as predicted by RNAfold (Figure S11) and 9027: G>A is a missense mutation that changes the codon to serine from glycine of mt-Co3 (22). It is interesting that both these SNVs are linked to glycine especially since glycine is an important part of biochemical pathways (glycine synthesis and metabolism, glutathione synthesis, glutamate-glycine cycle) in astrocytes (26) and also can act as an inhibitory neurotransmitter in neurons (51). It is likely that there are differences in the tRNA pools availability for neurons vs astrocytes. The 9027: G>A in mt-Co3 (an important component of OXPHOS) observed more frequently in neurons may suggest some advantage to mitochondria since neurons are known to critically depend on OXPHOS for energy. Moreover tRNA for serine requirement (due to glycine to serine change) is met more easily in neurons than in astrocytes. These findings suggest that the same mt-SNV sites may undergo different mechanisms of development, maintenance, and/or selection. Given that postnatal wildtype cells exhibit these differences, it suggests that the distinctive functions of mitochondrial bioenergetics may inherently be reflected at the mtDNA level and may potentially become pronounced during disease pathology or aging processes. Mean allele frequency differences for other SNVs were notable, but significance was lost after corrections for multiple tests.

Co-segregation in mt-DNA has been reported in human patient samples and trans-mitochondrial cybrids (created by fusion of enucleated cell with another mt-DNA depleted cell with a nucleus to study specific interaction between the nucleus and mt-genome) during aging, onset of deafness and mt myopathy (52–54). While a variant by itself may not be able to elicit a bioenergetic change, together with other mutations these SNVs may contribute to inefficient mt functioning (53). Of the most prominent linkages in the wildtype C57BL/6 mouse strain was that of 6430: A>C with 6432: G>C (both sites within mt-Co1) resulting from a lack of recombination. The linkage at 9027: G>A in mt-Co3 with 9461: T>C in mt-Nd3,may suggest selective advantages due to plausible cis effects, which when disrupted lead to sub-optimal oxidative phosphorylation. Although there may be a mutational bias towards C<>T and G<>A substitutions that could potentially increase linkage, except for the 9027-9461 linkage, all other linkages are not transitions. For 9027-9461, we also analyzed a mouse oocytes Nanopore sequencing dataset (55), and found 9027:G>A and 9461:T>C were indeed on the same haplotype. Further, these findings suggest numerous SNVs that may be linked, and if so, this may be a consequence of selective advantages conferred at either the cis or trans level. Regardless of the mechanism, this implies that the influence of pathogenic or non-synonymous SNVs may be impacted by targeting their corresponding linked SNVs. Long read length sequencing on single-mtDNA will help assess the co-segregation status of sequences highlighted in the present study.

Previous studies analyzing mt-genomes from single cells as well as single mt-genomes from single cells have been limited by technical challenge in single mitochondrion isolation. Several findings pertaining to mt heteroplasmy have been at the organism (56), tissue (20), or at the cell level (55, 57, 58). However, hierarchical characterizations of mt variants at the single-mt level (22) are limited. In this study, we conducted an analysis of SNV from up to 50 individual mitochondria obtained from a single cell. Isolating homogeneous populations of mitochondria with specific characteristics is challenging due to this inherent heterogeneity. As we have examined multiple mitochondria but only a small fraction of the total mitochondrial population in a cell, it is probable that our analysis has targeted a subset of the population. This analysis was not limited by the isolation yield, but rather by the manual processing of single-mitochondrion samples (mtDNA amplification) on the day of isolation. However, in the future to increase the total throughput through single mitochondrion isolation and amplification method reported here, automatic liquid and a single cell single mitochondrion barcoding scheme will be developed and implemented.

## Conclusion

Despite limitations arising from the small number of analyzed cells per animal, these data provide valuable insights into variation in the single cell mitochondrial DNA SNV landscape. The single cell-single mitochondrion approach 1645 individual mitochondria) in our study reveals higher number of SNV sites per mitochondrion than previously reported attributable to increased experimental resolution. The single-mt SNV landscape suggests that different loci within the mt- genome are subject to different mechanisms of clonal expansion and segregation. The purification selection in mouse neurons and with some of the cell type differences being modest, it will be interesting to determine whether these differences are present and expanded during human neurological disease development. These data are fundamental to the understanding of how different variants can exhibit site-specific differential patterns of clonal expansion and inter- and intra- cellular variation, which narrows the knowledge gap in understanding the functional role of heteroplasmy in disease processes.

## Materials and Methods

### Cell culture

Based on the protocol outlined by Kaech and Banker, 2006 (59), some modifications were made for culturing primary cortical neuronal co-cultures. Briefly, the postnatal day 1 mouse pups (C57BL/6) were dissected to carefully isolate the cortex in cold Hank’s Balanced salt solution without Ca^2+^, Mg^2+^. The cells were then dissociated with 0.5% trypsin/EDTA in PBS followed by pipetting, trituration and passing the cell suspension through a cell strainer. Subsequently, the cells were plated on poly-D-lysine/laminin coated 12-mm coverslips in 35 mm plate after live cell counting with trypan blue to achieve plating density of 200,000 cells/ml. Co-cultures of neurons and astrocytes were maintained in MEM with B27 supplement and 1% FBS at 37°C in 5% CO_2_ incubator as previously mentioned in Morris *et al*. (2017). (22).

### Staining and Isolation of single cells

Dispersed mouse cortical neurons and astrocytes were seeded on the coverslips in a 35mm dish were stained with 100 nM of MitoTracker Deep Red FM (Invitrogen, M22426) at 37°C for 20 minutes in an incubator (5% CO_2_) at DIV (days in vitro) 3, 7, or 21 days. The cell media with MitoTracker Deep Red was replaced with fresh media and incubated for another 20 minutes. A single neuron or astrocyte was harvested using a micropipette from a coverslip based on morphology (see SI for details and representative images Figure S14) with MitoTracker labeled cells under the microscope (40X magnification, Olympus IX70), and then was collected in the 1.7 mL Eppendorf tube with 4 μl of cell lysis buffer. After spinning down briefly, the Eppendorf tube was incubated on ice for 10 minutes followed by addition of 15 μl mt buffer (220 mM sucrose, 68 mM mannitol, 10 mM KCl, 5 mM KH_2_PO4, 2 mM MgCl_2_, 0.5 mM EGTA, PH 7.2; 0.01% bovine serum albumin and 1X Protease inhibitor cocktail) to the Eppendorf tube. The single neuron or astrocyte lysates were then used for preparation of the flow cytometry sorting samples.

### Conjugation of anti-TOMM20 antibody on microbeads

The conjugation of anti-TOMM20 antibody (Recombinant Anti-TOMM20 antibody [EPR15581-39] - BSA and Azide free, Abcam Cat# ab220822, RRID:AB_3097753))) to microbeads through an avidin-biotin bond was modified based on the commercial protocol (Spherotech Inc., Particle coating procedures - https://www.spherotech.com/technical%20notes/STN-1%20rev%20C.pdf). The anti-TOMM20 antibody was biotinylated via Miltenyi One-Step Antibody Biotinylation Kit (Miltenyi Biotec, 130-093-385) following the manufacture’s protocol. 10μg of anti-TOMM20 antibody was diluted to 100μg/mL by adding 1X PBS, followed by addition of the10μg of antibody to the well containing Miltenyi lyophilized biotinylation mix. After pipetting thoroughly, the mixture was incubated at room temperature for 24 hours and stored at 4°C before use. For coating microbeads, 2μg of biotinylated anti-TOMM20 antibody was suspended in sodium phosphate buffer (PB buffer, 0.1M, pH=5.5) at 10μg/mL.20 million avidin coated polystyrene particles (1.7-2.2 μm, 0.1% w/v, Spherotech Inc, VFP-2052-5) were added and incubated with biotinylated antibody for 1h at room temperature on rotator covered by aluminum foil. After incubation, the coated microbeads were centrifuged at 150,000g for 10 minutes and resuspended in 0.4mL PB buffer. The centrifugation of beads was repeated, and the beads were resuspended in 0.4mL of filtered mt buffer mentioned in the previous section. Anti-TOMM20 antibody coated beads were freshly prepared each time for mitochondria isolation.

### Flow cytometry of captured mitochondria from single cells

Anti-TOMM20 antibody conjugated fluorescent microbeads (1-2 million - 10 times of the estimated number of mitochondria from individual cells) were added to the single neuron or astrocyte cell lysate and the final volume of each sample was adjusted to 100μL by adding filtered mt buffer. After an hour of incubation at room temperature on a shaker, the product was transferred to flow tubes for FACS sorting. Flow cytometry was performed on a LSR II flow cytometer (BD Biosciences) and BD FACS Aria II (BD Biosciences) respectively for assay validation and sorting at the Penn Cytomics and Cell Sorting Shared Resource Laboratory (<u>RRID:SCR_022376</u>). Flow cytometric analysis was performed using FlowJo software (Figure S15).

### Amplification of mt-genomes by Rolling Circle Amplification (RCA)

Single mitochondria were sorted in each of the wells of a 96-well plate containing 4 μl of mild lysis buffer (1x TE, 0.1M NaCl, 2mM CaCl_2_). 1μl of 0.25 μg/μl Proteinase K (Roche, 03115879001) was added to each well and incubated at 55°C for 30 mins. The Proteinase K inactivation was performed by incubation at 95°C for 10 mins. The plates were rested on ice until the next step. Nuclease free water and 1X Phi 29 reaction buffer (New England Biolabs, B0269SVIAL) were added to each well followed by incubation at 70°C for 3 mins. The plates were immediately rested on ice for at least 2-4 mins to complete denaturation. The samples were incubated at 25°C for 50 minutes with random primers (20μM). 2mM dNTPs (Invitrogen, 18427088),10mM DTT (Thermo Scientifc,707265ML) and 5U Phi 29 DNA polymerase (New England Biolabs, M0269L) were added to each well to set up the reaction. The 25 μl reaction mixture was incubated at 30°C for 32 hours. Another 5U of Phi 29 DNA polymerase was added to the reaction mixture after 20 hours to continue the reaction. The reaction was stopped after 32 hours by incubation at 65°C for 10 mins. The RCA product was stored at -20°C until the next step. The amplified mt-genome was validated by ethidium bromide agarose gel electrophoresis (Figure S16).

### Library preparation

For each library, 10 Non-Seq Barcodes (namely, M1 through M10, Table S1) were designed for multiplexing 10 mitochondria. For each mitochondrion, 12 target regions were further PCR amplified using forward primers beginning with the aforementioned Non-Seq Barcodes and reverse primers as listed in Table S2. These regions were selected based on the SNV sites reported in our previous paper (22). All primers were designed through the NIH Primer-Blast tool (https://www.ncbi.nlm.nih.gov/tools/primer-blast) and synthesized by Integrated DNA Technologies. PCR amplification was performed with 1 μl of RCA product from each single mitochondrion sample, 1μl of 24 oligos pool for 12 target region amplification (1.25μM), 23 μl Platinum PCR SuperMix (Invitrogen, 12532-016), using the following program: 94°C 2 min for the 1^st^ cycle, followed by 25 cycles of 94°C 30s, 49°C 30 s, 68°C 30 s, followed by addition of 15 μl Platinum PCR SuperMix for next 20 cycles using the same PCR program as the previous 25 cycles, with a final hold at 4°C.

After the amplification, each PCR products were analyzed on the Agilent Type Station 4150 using a High sensitivity D5000 Screen Tape Assay. Then 10 barcoded single mitochondria were pooled together and cleaned up by 1.5X AMpure XP beads (Beckman Coulter, A63881). 50 ng of the pool was input into a library using TruSeq Nano DNA kit (Illumina, 20015964) as per the manufacturer’s instructions. Twenty-three libraries were combined into a sequencing run and then sequenced by 1x150bp single-end kit on Illumina NextSeq 500. For more details on sample size for the number of mitochondria isolated per cell from each mouse pup (Figure S1A, S17) and PCR read depth (Figure S18), see SI.

### SNV calling within each mitochondrion sample and Quality Control Procedure

We processed the raw reads using an in-house Next-Generation Sequencing (NGS) pipeline. First, barcoded reads from the same library were demultiplexed into individual mt reads. Then, sequencing adapters and poly-Gs were searched, trimmed, and reads shorter than 20bp were discarded. Remaining reads were mapped to the mouse mtDNA (genome build mm10) using STAR,v2.7.2d (RRID:SCR_004463) (60) with default parameters except ‘-- outFilterMismatchNoverLmax 0.1’, which made mapping more stringent than by default. To disable spliced alignment, additional arguments ‘--alignIntronMax 1 --alignSJDBoverhangMin 999’ were used. Afterwards, only uniquely mapped reads inside the PCR regions were extracted, and for each PCR region, per-base read pileup was generated by samtools, v1.13 (RRID:SCR_002105) with parameters ‘--excl-flags UNMAP,SECONDARY,QCFAIL --count- orphans --min-BQ 0 --min-MQ 0 --reverse-del’. In case of memory exhaustion, an additional argument ‘--max-depth 500000’ was used such that the maximum depth was limited to 500,000. To eliminate sequencing errors, bases with Phred scores < 30 were excluded. The details regarding read alignment and cutoff choice can be found in the SI (Figure S19). For accurate AF estimation, SNVs were called from positions with (1) read depth ≥ 50, and (2) VAF ≥ 5% (Dataset S8). The PCR primer regions were discarded from the SNV calls. To reduce false positives, we discarded PCR primer regions from SNV calls or SNVs discordant between the alignment tool STAR (60) and BWA (RRID:SCR_010910) (61) (Figure S20). Besides the original SNV call, another much more stringent SNV list was derived by the following filtering criteria: (1) reads must be on the positive strand only, (2) reads must start off the 5’ end of the PCR forward primer only by +/-1 bp, and (3) read length must be >= 135bp. The stricter SNV call verified 92.5% of the total SNVs from the original call. As compared to negative controls, single-mt samples showed significantly higher read depth and positive controls (pooled mtDNA from mouse brain) showed the highest depth. The quality assessment of positive and negative controls (Figure S21) as well as the fix to variant 9027:G>A that was lost due to soft-clipping (Figure S22) can be found in SI and Dataset S9. Given our single cell fluorescence activated mt isolation procedure (23), we do not expect nuclear DNA contamination in our samples. In support of this argument that nuclear-embedded mitochondrial DNA Sequences (NUMTs) (62–65) are highly unlikely in our dataset (Figure S23 and Dataset S10), please refer to SI for computational prediction of NUMTs.

### SNV impact annotation

The package Variant Effect Predictor, VEP - v102.0 (RRID:SCR_007931) (66) with the parameters ‘--offline --cache --species mus_musculus -- force_overwrite --everything -i’ was used to infer the consequence of the 1032 non-reference alleles based on the mouse mt codons. We manually labelled AUU-to-AUC (e.g. at 9461) as ‘synonymous’ according to the mouse mt codon table. Besides, 3772:C>T hit both mt-Ti (on the plus strand) and mt-Tq (on the minus strand) while VEP predicted identical severity; only former was retained for simplicity.

### Comparing SMITO and sites segregating amongst mouse strains

To assess the data quality SI, Figure S24), 17 mouse strains’ mt-genomes data from the Sanger Mouse Project (67, 68) were aligned (multiple sequence alignment) using Clustal Omega (RRID:SCR_001591) (69), which identified 1494 segregating sites. Among them, 128 sites within the 12 mt target regions analyzed in our study were termed as ’17 mouse strains data’ and compared to SMITO dataset. The overlap was quantified by hypergeometric tests, and by major allele or top minor allele consistency.

### Inherited-somatic SNV demarcation

To compute the null distribution of number of SNVs shared by *k* mice with animal specificity controlled, the SNVs in each mouse were shuffled such that the total number of SNVs remained identical in each permutation (*n* =100), and the average was calculated as the expected number of SNVs shared by *k* mice. The expected and the observed curve were contrasted; the first cross (*k* <1.9) was taken as the cutoff for somatic SNVs; inherited SNVs were taken only if *k* ≥3 for stringency.

### Mutational pattern in annotation groups

To compare mutation pattern for somatic and inherited SNVs, the total SNVs were divided into two groups: coding and non-coding.. In each group, the per-base mutational rate and the transition-to-transversion (Ti/Tv) ratio was compared using Fisher’s exact test (two-sided)

### Hierarchical variation distribution of the inherited SNVs

ANOVA was performed on the VAF of 161 inherited SNVs and the Mean of Squares (MS) was computed at three levels: the mouse, cell, and mitochondria. For each SNV, the ratio of the mouse-level MS to cell-level MS, and of the cell-level MS to mitochondria level MS was calculated. To ensure that the distribution of the MS was not skewed by noise (VAF < 5%), mouse and cell labels were randomized by 1000 permutations, creating a null distribution of the mouse-, cell- and mitochondria-level MS for the observed distribution to be contrasted. For each SNV, the MS at three levels were finally normalized to a percentage. This analysis involves cell-level and mouse-level variance partition where mouse identity has to be ascertained, so the mixed pup culture (16&17) was excluded.

### Cell type comparison for SNV incidence and VAF variance

The number of SNVs per mitochondrion (log(x+1) transformed) were compared in neurons and astrocytes by a mixed-effects linear model using the cell type as the main effect, the mouse ID as the random effect, and the number of bases with sufficient depth (log(x+1) transformed) as covariates; the null distribution was built by 1000 permutations to the cell type label. Each SNV incidence between the cell types were compared using two independent methods: (1) logistic regression was performed to predict the presence or absence of an SNV in individual mitochondria using the cell type as the main effect and the mouse ID as the random effect; (2) the fraction of mitochondria with variant in each cell (excluding cells with < 3 mitochondria) was calculated and then fitted to a weighted logistic regression using the cell type as the main effect and the mouse ID as the random effect. R package lme4,v1.1-28 lme4 (RRID:SCR_015654) (70) was employed for the mixed-effects models. Finally, the between-mitochondria variance between the cell types for each SNV was compared. To control for the mouse and the cell effect, the VAF was arcsine transformed and ANOVA was performed using the mouse and cell ID as the covariates, and the MS of the model residuals from the cell level was taken as the between-mitochondria variance estimate. The analysis was performed for both cell types separately. Mouse16&17 is a co-cultured sample from two mice (mixed pup culture), consequently cells from this experiment were excluded from the analyses in Figure 5. The method by Li *et al*., 2015 (21) was used to compute Ka/Ks statistic controlling for the mutational spectra. The null distribution was built by 3000 permutations controlling the respective mutational spectrum in either cell type.

### SNV linkage analysis

To detect linked SNVs, the cells that had ≥ 20 mitochondria were retained, followed by filtering for the SNV sites that had non-missing allele calls in ≥10 mitochondria and that had ≥5% mitochondria with the minor allele (SI, Figure S25-26). The association between a pair of SNVs was quantified by a one-tail Fisher’s exact test. Briefly, for each cell and for each pair of the filtered SNV sites, whether the observed odds ratio of any combination of observed alleles significantly exceeded the random was tested. The *p*-values were subsequently corrected for multiple tests using the Benjamini-Hochberg method (71).

#### Statistics

Statistical analyses and graphics are done with R Project for Statistical Computing (RRID:SCR_001905), circlize (RRID:SCR_002141) and ggplot2 (RRID:SCR_014601).

## Declarations

### Ethics approval and consent to participate

Not applicable

### Consent for publication

Not applicable

### Availability of data

The raw sequencing reads, the metadata and the processed AF table have been deposited to GEO under the accession number GSE218122. The data preprocessing and SNV calling pipeline are available at https://github.com/kimpenn/smito-pipeline. Scripts for data analysis and figures are publicly available at https://github.com/kimpenn/smito-analysis. Novel experimental reagents are available upon request.

### Authors’ contributions

Conception and design of the experiments: P.S.K., Z.Y., H.Z., Y.L., D.I., and J.E., collection of data: P.S.K., Z.Y., H.Z., Y.A., and N.S., analysis of data: Y.L., J.K., E.N., S.F., P.S.K., Z.Y.,, D.I., and J.E., interpretation of data: Y.L., J.K., P.S.K., Z.Y., D.I., and J.E., drafting and revising the article critically: P.S.K., Z.Y., Y.L., J.K., D.I., and J.E.

## Acknowledgements

The authors would like to thank Kevin Miyashiro for preparing primary mouse cultures, Shifu Tian for assisting at the Flow Cytometry and Cell Sorting Facility, and Zhen Miao for statistical discussion.

## Author Information

Parnika S Kadam, Zijian Yang, Youtao Lu, and Hua Zhu contributed equally to this work. Junhyong Kim, David Issadore, and James Eberwine contributed equally to this work. Correspondence to David Issadore and James Eberwine.

