## Supplementary Information for "Single-Mitochondrion Sequencing Uncovers Distinct Mutational Patterns and Heteroplasmy Landscape in Mouse Astrocytes and Neurons"

**Supplemental Methods.**

**Morphology criteria for single cell pickup:** Since both neurons and astrocytes can show variable morphologies, we used the following criterion for each cell type.

Neuron – Pyramidal neurons (A), bipolar (B) or unipolar (C), were selected (Figure S14 A-C). A cell was classified as a neuron if it had a small soma (approximately <10 um) with elongated 1-3 processes.

Astrocytes – Cells displaying 'classic-star shaped' morphology with multiple processes were chosen as astrocytes. A cell with a relatively larger (>10 um) and flatter cell body, accompanied by more than 4 processes of similar lengths were designated as astrocytes (Figure S14 D-F).

Additionally, we have incorporated references demonstrating that the neurons and astrocytes chosen for analysis exhibited morphological characteristics consistent with those of mouse cortical neurons and astrocytes identified by immunofluorescent markers. Please refer to the following references showing neuronal and astrocyte morphology that resembles the cells we have selected as mouse cortical neurons and astrocytes respectively: Hilgenberg and Smith 2007 (1), Sciarretta and Minichiello 2010 (2), Granger et al. 2020 (3), Xu et al. 2022 (4), Baldwin et al. 2023 (5), Sathe et al. 2017 (6), Sosunov et al. 2020 (7), Sun and Jakobs 2012 (8), Purvis et al. 2022 (9), Sidoryk-Wegrzynowicz et al. 2017 (10), Sun et al. 2017 (11), to name a few.

**Mitochondrial (mt) barcode demultiplexing.** To determine the optimal threshold of error tolerance in mt barcode demultiplexing, the Levenshtein distance was calculated between any two barcodes (Table S1B) and a minimal distance 3 was obtained. Therefore, at most two errors in searching each of the 11bp barcodes were permitted using the command: cutadapt -g XCGATTATCACG -e 0.2 -O 11 (CGATTATCACG being M1 as an example,

Table S1A). Please refer Figure S17-18 for sample overview and PCR read depth for each mt barcode.

**Out-of-range SNV filtering.** The mt barcode and the PCR primers' efficiency by the read depth was examined at each base position. We noticed the trailing coverage at the 3' end of some PCR target regions (e.g., Region3, 6 and 7, Figure S18). For the sake of rigor, the bases out of the sequencing read length were designated as "out-of-range", SNVs (if any) inside these regions were discarded. In addition, SNVs (if any) inside the PCR forward primers were removed because the genetic variants could be masked by the identically synthesized primer sequences, and hence any SNVs inside the primers could be artifactual.

**Base Phred score filtering.** To prevent false positives caused by sequencing errors, polymorphic bases with Phred scores below 30 were discarded. This threshold was derived from the Phred calibration analysis, where all non-reference bases were collected and their theoretical (x-axis) error rate was plotted from Phred calls against the observed error rate (y-axis), which was composed of true SNVs and technical errors. As expected, two well separated clusters were observed: the top-right quadrant (VAF >= 5%, Phred >= 30) and the bottom-left quadrant (VAF < 5%, Phred <30) (Figure S19). The threshold was maintained at 30 for the sake of simplicity (not lowered to 28.99 -the red dashed line). Only < 10% read depth was lost when imposing the more stringent cutoff, and most of the loss occurred to the 3' end.

**Variant allele frequency threshold.** Assuming that the RCA reaction went through $n_1$ cycles at a per-base error rate $\varepsilon_1$, and that the PCR reaction went through $n_2$ cycles at a per-base error rate $\varepsilon_2$. For a particular site, let $X_1$ be the total number of mutations resulting from RCA, and that $X_1 \sim \text{binom}(n_1, \varepsilon_1)$, hence after RCA $X_1$ mutant copies and $n_1 - X_1$ intact copies will be obtained. Assuming no backward mutation, all mutated molecules would be amplified by PCR, and the final number of mutant copies would be $X_1 \times 2^{n_2}$. For each of the RCA intact copies, assuming it induces $X_2^i$ errors during PCR cycle $i$, where there are $2^{i-1}$ reactions, hence $X_2^i \sim \text{binom}(2^{i-1}, \varepsilon_2)$, and these errors would propagate and eventually result in $2^{n_2-i} \times X_2^i$ mutant copies. Therefore, the total number of mutant copies exclusive to PCR would be $(n_1 - X_1)\sum_{i=1}^{n_2} 2^{n_2-i} \times X_2^i$. After RCA and PCR, the total number of amplicons would be $N = n_1 \times 2^{n_2}$ and among them, the total number of mutants would be $X = X_1 \times 2^{n_2} + (n_1 - X_1)\sum_{i=1}^{n_2} 2^{n_2-i} \times X_2^i$. Assuming the amplicons to the depth of $M$ were sequenced and $Y$ variant bases were observed, then $Y \sim \text{binom}\left(M, \frac{X}{N}\right)$, hence at the VAF threshold $\theta$, the probability of making a false positive call is $\Pr(Y > M\theta)$.

For simplifying $\frac{X}{N}$.

$$\frac{X}{N} = \frac{X_1 \times 2^{n_2} + (n_1 - X_1)\sum_{i=1}^{n_2} 2^{n_2-i} \times X_2^i}{n_1 \times 2^{n_2}} = \frac{X_1}{n_1} + \left(1 - \frac{X_1}{n_1}\right)\sum_{i=1}^{n_2} 2^{-i} \times X_2^i$$

Assuming $X_1$ and $X_2$ are independent random variables, we further have

$$\mathrm{E}\left(\frac{X}{N}\right) = \mathrm{E}\left(\frac{X_1}{n_1}\right) + \mathrm{E}\left(1 - \frac{X_1}{n_1}\right)\mathrm{E}\left(\sum_{i=1}^{n_2} 2^{-i} \times X_2^i\right) = \varepsilon_1 + (1 - \varepsilon_1)\sum_{i=1}^{n_2} 2^{-i} \times 2^{i-1} \times \varepsilon_2$$

$$= \varepsilon_1 + \frac{1}{2}n_2(1 - \varepsilon_1)\varepsilon_2 \approx \varepsilon_1 + \frac{1}{2}n_2\varepsilon_2$$

Given that $\varepsilon_1 = 5 \times 10^{-6}$, $\varepsilon_2 = 3.67 \times 10^{-6}$, $n_2 = 45$, we have $\mathrm{E}\left(\frac{X}{N}\right) \approx 8.758 \times 10^{-5}$. Since the post-filtering minimal sequencing depth $M = 50$ is sufficiently large and $\mathrm{E}\left(\frac{X}{N}\right)$ is sufficiently small, we can approximate $Y$ with a Poisson distribution with $\lambda = 4.379 \times 10^{-3}$.

Therefore, at the VAF cutoff $\theta = 1\%, 2\%, 5\%, 10\%$, the false positive rate to be $\Pr(Y > M\theta) = 4.370 \times 10^{-3}, 9.560 \times 10^{-6}, 1.395 \times 10^{-8}, 9.756 \times 10^{-18}$, was expected respectively. Given 1424 sites in total, the expected false positive sites were $6.222, 1.36\ \times 10^{-2}, 0, \text{and}\ 0$ at the corresponding cutoff. Hence 5% VAF threshold was chosen.

**Control experiment assessment.** The raw read depth in the single mtDNA samples versus different types of controls was compared: pooled mtDNA(+), pooled mtDNA(+) primer(-), pooled mtDNA(-) primer(+), pooled mtDNA(-)primer(-) in 12 PCR target regions (Figure S21). As expected, in most of the amplified SNV regions, the positive controls (i.e., pooled, double positive) gave rise to the highest read depth, and the single-mitochondrion, double positive samples often showed the second highest depth. Compared with the positive controls, the negative controls generally showed two orders of magnitude lower depth. It is noteworthy that the read depth is not an accurate estimator for sensitivity for it is heavily influenced by PCR efficiency, starting material and sequencing depth, hence single-mitochondria samples and pooled controls might not be well comparable. The main purpose of this analysis was to highlight the global difference between the positive and negative pooled controls, and the results did match our expectation.

**Correction for the soft-clipped 9027.** During read alignment, we noticed that STAR could soft-clip the mismatching base(s) at the 3' end of reads and hence could cause an underestimated VAF should the variant base be at the 3' end. A systematic diagnostic analysis was performed to evaluate the impact of this issue. We found that 9027 was the only position affected by this problem—of 1489 mitochondria that have base coverage at 9027 from raw reads, there were in total 107,259,677 As, 13,953,942 Gs, 330,682 Cs, and 294,080 Ts; STAR discarded the non-reference As. Therefore the 9027:G>A read frequency for each sample was manually compensated. Figure S22 represents the change in 9027:G>A VAF before and after the correction.

**Mouse strains coordinate standardization.** During the phylogenetic analysis, a subtle discrepancy was seen in the UCSC GRCm38 reference genome (C57BL/6J) which is used in this study and the C57BL/6NJ genome (CM004277.1) from the Mouse Genomes Project: the former has 16299bp while the latter has 16300bp (Figure S24). Hence a distinction was made between C57BL/6J and C57BL/6NJ: UCSC C57BL/6J was added to the 16 strains from the Mouse Genomes Project and multiple sequence alignment was performed to 17 strains in total. The mitochondrial coordinate system was standardized based on UCSC GRCm38 (C57BL/6J).

**Alignment quality assessment for linkage analysis.** During analyzing SNV linkages, some significantly linked SNVs were observed in a small fraction of amplicons and showed suboptimal read quality. For example, 6 variants around 13775 were located on reads starting ~80bp downstream the 5' end of the PCR forward primer (Figure S25) and appeared as linked SNVs. To avoid suboptimal alignment that could be a major confounder for this analysis, a stricter set of filtering criteria was used: (1) reads must be on the positive strand only, (2) reads must start off-5'end only by +/-1 bp, (3) read length must be >= 135bp. Using this stricter processing, 776 total SNV sites and 209 SNV sites shared by >=3 mitochondria were observed. Comparing with the permissive processing (Figure S26), 92.6% (776) SNV sites survived the stricter filtering (Figure 26A). The stricter filtering also led to 22 new SNV sites shared by >= 3 mitochondria because of an increase in the VAF after the stricter filtering (Figure S26B).

**NUMTs Computational Analysis on SMITO data.** To determine the number of false positives due to the potential NUMTs (12–16) contamination, we first assembled a list of mouse NUMTs regions. Briefly, we synthesized 50 bp reads at each mitochondrial position and mapped them to the nuclear genome using BLAST with highly sensitive parameters: -num_alignments 1000 -word_size 6 -perc_identity 80 -gapopen 3 -gapextend 1 -evalue 1; the results were merged as disjoint regions and referred to as NUMTs. We synthesized 50 bp reads from NUMTs and mapped them back to the mt-genome using bwa mem. To make the mapping quality comparable to STAR, we filtered for alignments with MAPQ >= 60. Finally, we called 381 NUMTs original, false mt-SNVs (Supplementary Fig S23A, Supplementary Dataset S10) using the aforementioned procedure in Methods section. Please note that, for this analysis we only required VAF > 0 and did not filter out low-depth or low-VAF SNVs, hence our false positives here are overestimated. We compared our SNVs against this list; only 27 overlapped suggesting that, by computational analysis at most, 27 out of 838 (3.2%) SNVs could potentially be artifacts originating from NUMTs.

Furthermore, mapping reads that did not align to the mouse mt-genome to the mouse nuclear genome yielded a minimal number of unique alignments (average 0.33%, median 0.08%). Given the same amplification, library prep and sequencing conditions, nucleus borne reads should have mismatch rates comparable to mt reads. These nucleus exclusive alignments showed significantly (Wilcoxon rank-sum test $p$-value <2.2e-16) higher mismatch rate (mean 5.19%, median 5.27%) than mt exclusive alignments (mean 1.41%, median 1.17%).

**PCR to target regions in nuclear genes.** To check if our single mitochondrion samples are contaminated with nuclear material, PCR was performed on the single mitochondrion RCA products using primers targeting a region from mouse mt-genome (Target region 6) and two independent regions in mouse chromosome 1 (Pet-1, FEV transcription factor) and chromosome 6 (Gapdh, Glyceraldehyde-3-phosphate dehydrogenase). Both 1µl and diluted single mitochondria RCA product (2µl of 1:25 dilution) was used to test the presence of mtDNA as well as nuclear contamination following the procedure mentioned in the main methods section under library prep. The annealing temperatures of 50°C, 55°C, and 57°C were used respectively for target region 6, Pet-1, and Gapdh. The isolated mtDNA (0.1pg) from mouse pup brain that had also undergone RCA was used as a positive control template (0.5µl) for target region 6 and 12 ng of mouse genomic DNA from mouse pup brain was used as template for positive controls, Pet-1 and Gapdh. 1µl from final PCR mix of test samples and 0.5µl of positive controls was loaded on Agilent D5000 ScreenTape. Fig S23B is a representative image showing PCR product from 4 single mt samples (from two independent experiments #45 and #22 – well IDs are labelled on the gel picture H1, H2 from #45 and B3, B4 from #22) diluted 1_25 samples targeted for target region 6 (Lane 2-5), Pet-1(Lane 6-9), Gapdh (Lane 10-13), positive control for target region 6(+Mt – Lane 14), positive control for Pet-1(+Nu – Lane 15), and positive control for Gapdh (+Nu – Lane 16). The expected band for target region 6 is 189 bp, Pet-1 is 237 bp and Gapdh is 452 bp for mRNA and 646 bp for DNA. The primer information for target region 6 is mentioned in Table S2. The primer information 5'-3' for Pet-1 and Gapdh are as follows:

Pet-1 Forward Primer – AGATTCTGGAACTCCCGTGT

Pet-1 Reverse Primer – GAGAAAGGGAAGCCAGAGTG

Gapdh Forward Primer – ACCACAGTCCATGCCATCAC

Gapdh Reverse Primer – TCCACCACCCTGTTGCTGTA

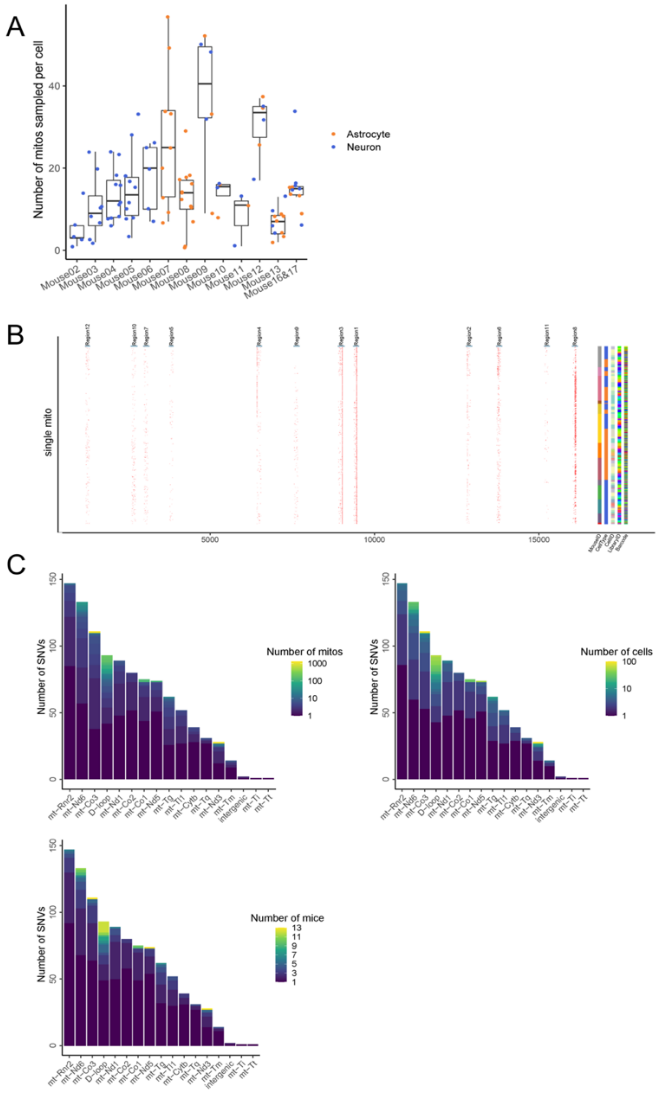


**Figure S1. An overview of the distribution of the total SNVs.** (A) The distribution of the number of mitochondria captured per cell per mouse. The mitochondria number isolated per neuron or astrocyte is depicted as a blue or an orange dot respectively. (B) The graph depicts a SNVs' presence (as red dot) in each mitochondrion sample on the y-axis (each row is one individual mitochondrion sample) across the mt-genome ( x-axis). The y axis on the right shows identity of each mitochondrion sample with respect to the mouse, cell type , cell, library ID barcode(C) The number of SNVs is represented on the y-axis in each target region genes depicted on the x-axis. SNVs color coded by the number of shared mitochondria, cells, and mice.

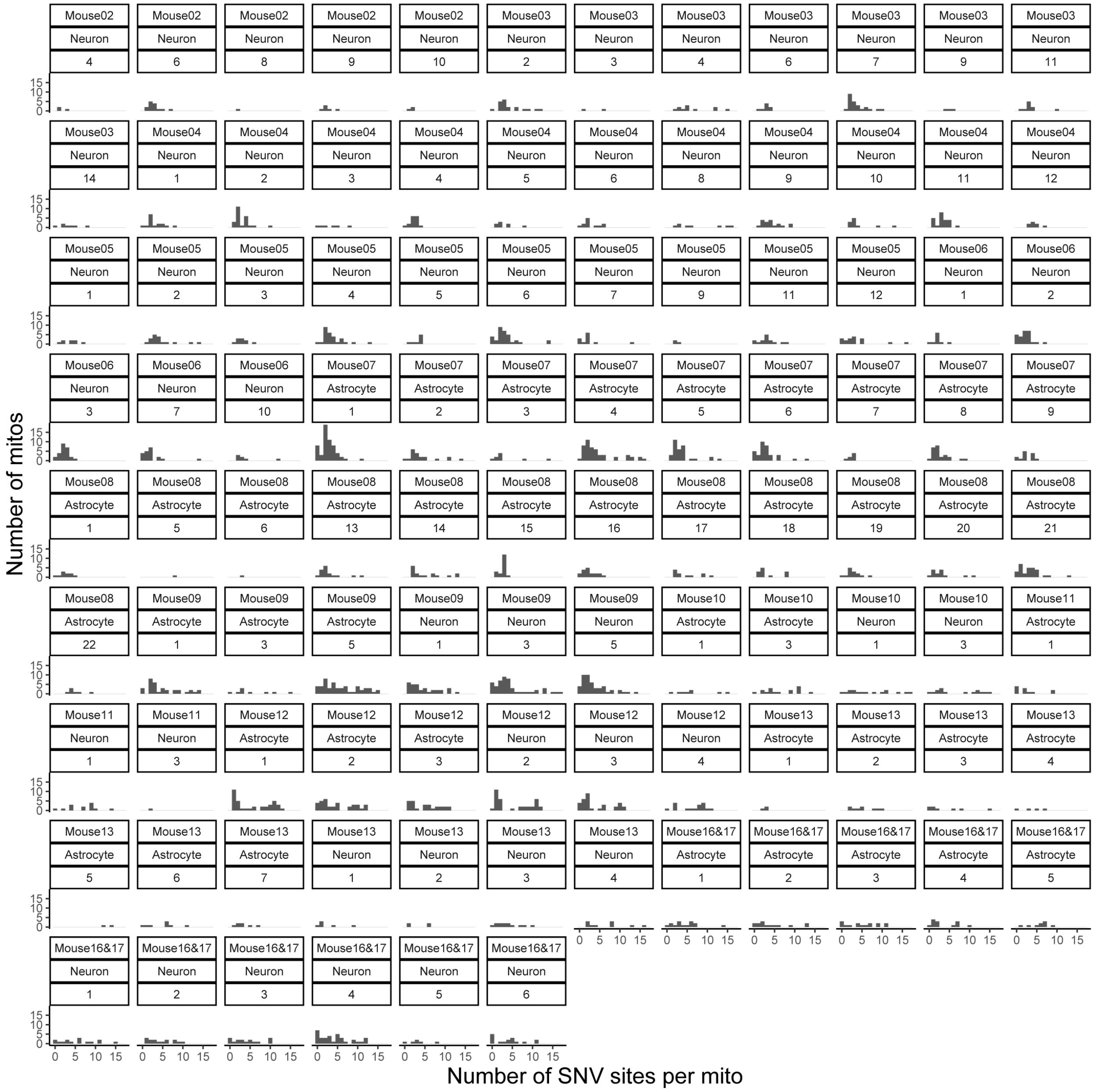


**Figure S2. Distribution of the SNV counts in the SMITO dataset.** Histograms showing the distribution of the SNV count in each mitochondrion for each of the 102 cells analyzed. Each histogram represents a cell (102 total) and is labelled with the cell type and the animal ID.

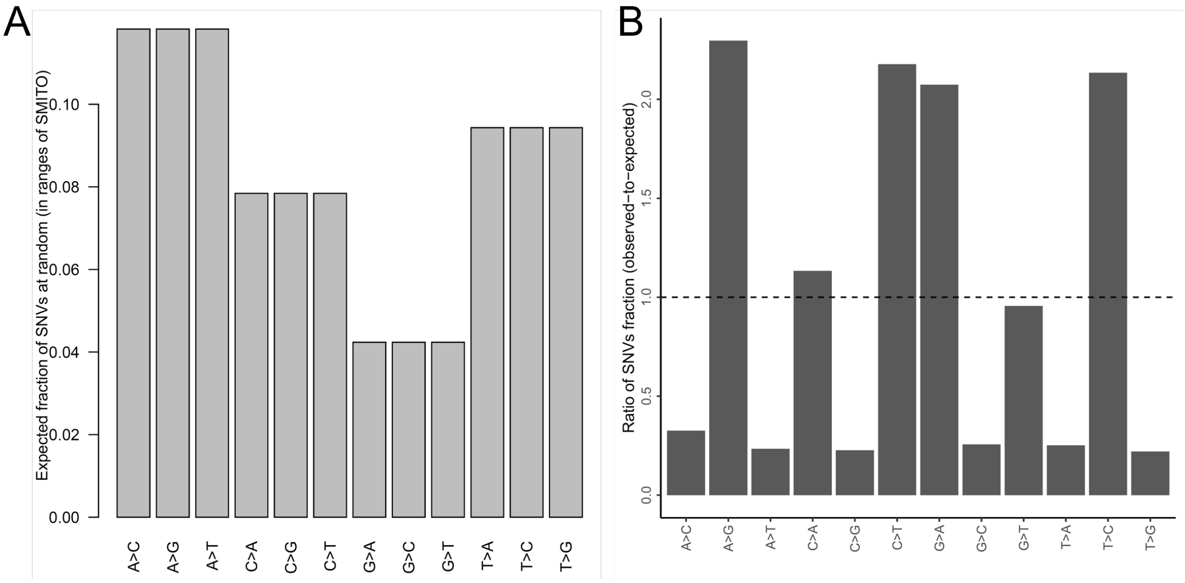


**Figure S3. Mutational Spectra for the mt-genome. (**A) Intrinsic C<>T, G<>A tendency due to mt-genome base composition and (B) the SMITO mutational spectra accounting for the intrinsic tendency.

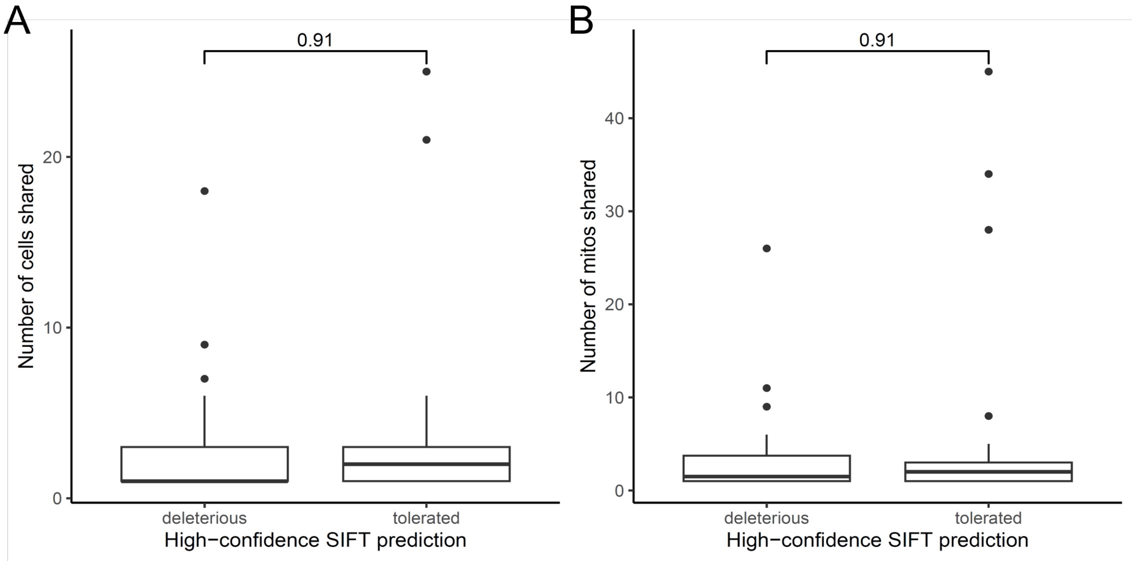


**Figure S4. Comparison of Cells and Mitochondria Sharing Deleterious and Tolerated Nonsynonymous SNVs.** (A) Number of cells or (B) mitochondria sharing the nonsynonymous SNVs, which were categorized as “deleterious” and “tolerated” by SIFT (for high-confidence only). The *p*-value is from Wilcoxon’s rank-sum test.

The Ac-loop of mt-Tg (5' -> 3')

```
-2-101 23
CT[TCC]AA  becomes
CT[TCT]AA
```

**Figure S5. Illustration of the anti-codon base changes in a tRNA SNV 9423:C>T.**

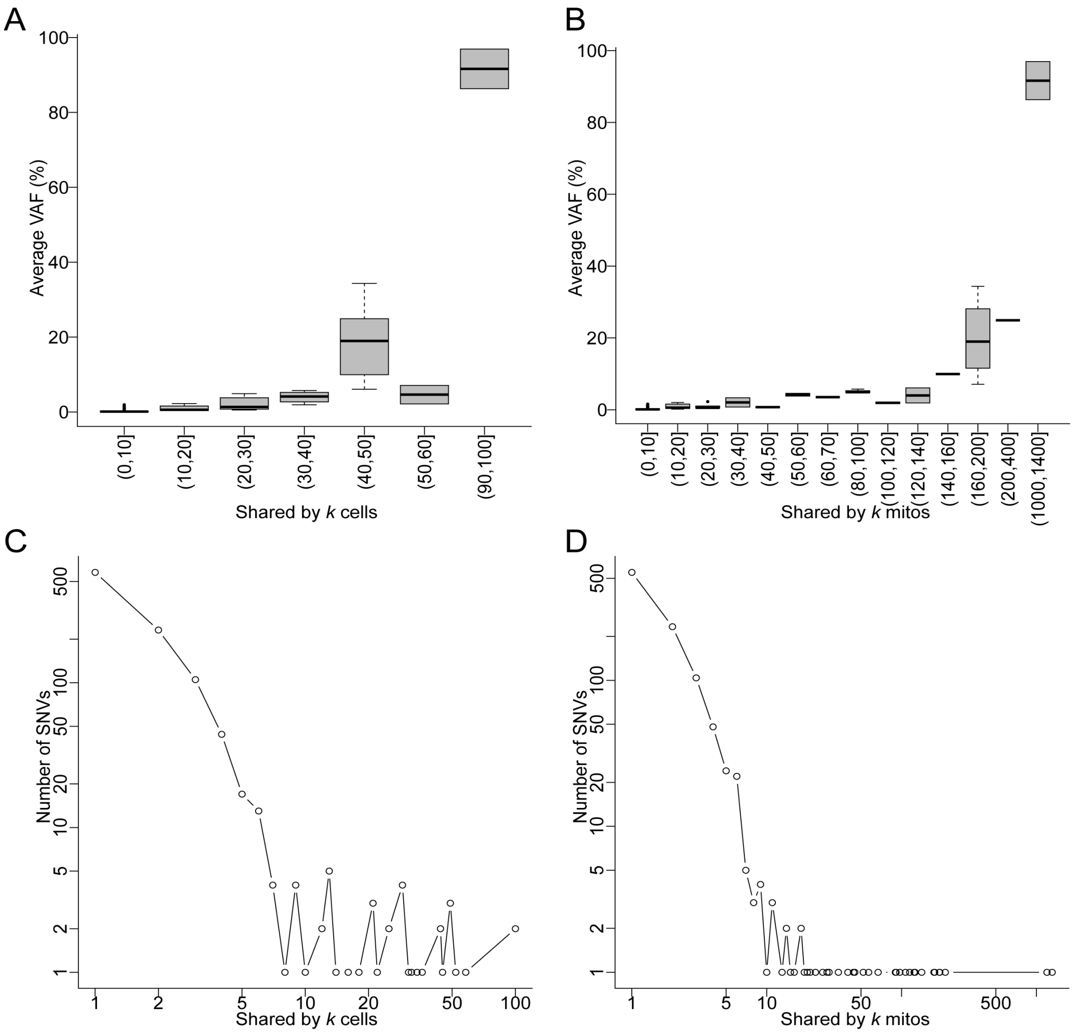


**Figure S6. The distribution of the VAF and number of SNVs shared by the single-mt samples.** (A) The relationship between the average AF (y-axis) and the number of cells sharing the SNVs. (B) The relationship between the average AF (y-axis) and the number of mitochondria sharing the SNVs. (C) The number of SNVs (y-axis) stratified by the number of cells sharing these SNVs. (D) The number of SNVs (y-axis) stratified by the number of shared mitochondria with these SNVs.

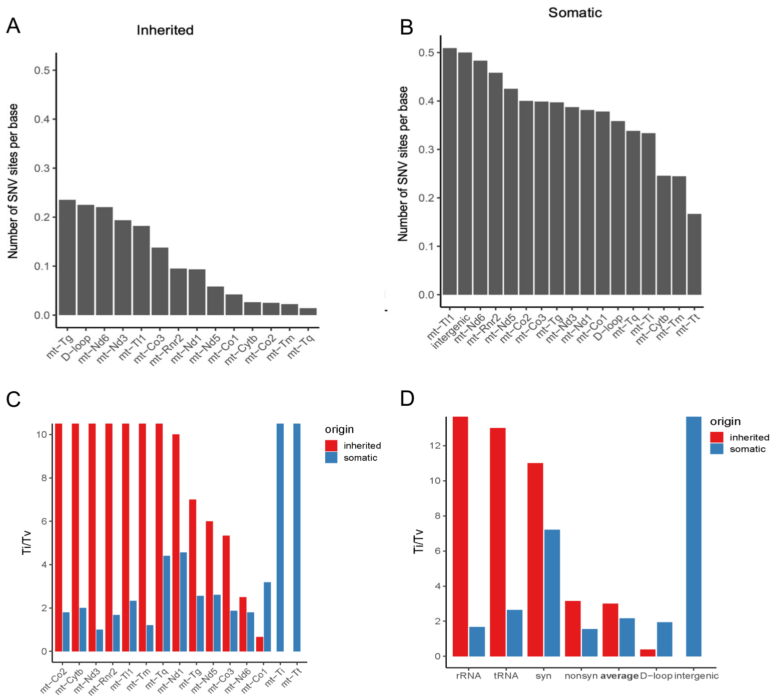


**Figure S7. Comparison between the inherited and the somatic SNVs in mutational propensity.** (A) Inherited SNVs' per-base variant rate on the y-axis in each target gene loci on the x-axis. (B) Somatic SNVs' per-base variant rate on the y-axis in each target gene locus on the x-axis. (C) The Ti/Tv ratio of inherited (red) and somatic (blue) SNVs for each target region gene. (D) The Ti/Tv ratio of inherited (red) and somatic (blue) SNVs for each group namely rRNA, tRNA, synonymous, nonsynonymous, D-loop, intergenic and the overall average). In (C-D), highest bars were capped due to infinite ratios (no transversion).

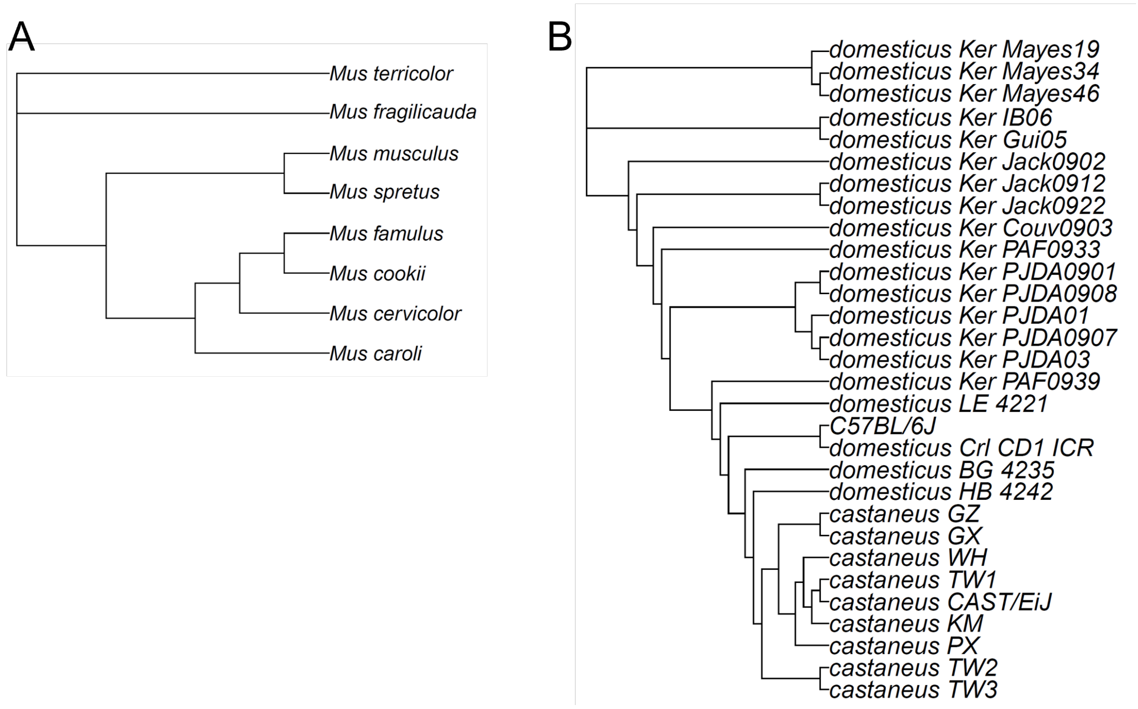


**Figure S8. Neighbor-joining phylogenetic trees.** (A) Phylogenetic trees of 8 Mus species and (B) two Mus musculus populations.

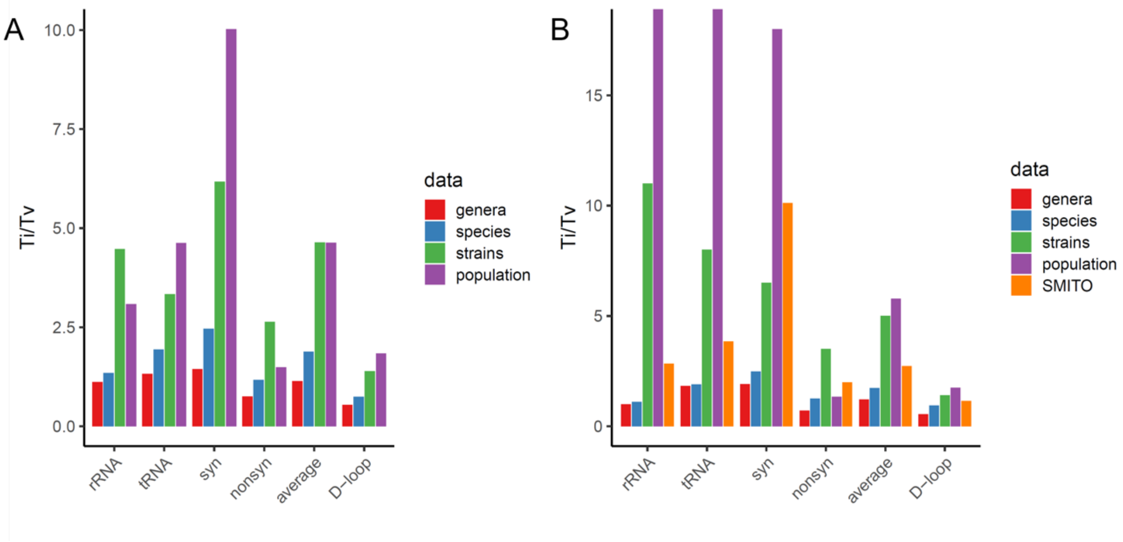


**Figure S9. Ti/Tv ratios for segregating sites** (A) whole mt-genome, (B) and regions assayed by SMITO between (1) genera (Mus musculus and Rattus norvegicus), (2) Mus species, (3) Mus musculus strains, (4) Mus musculus populations in respective functional class. In panel B, the highest bars were capped due to being infinite (i.e. transitions only).

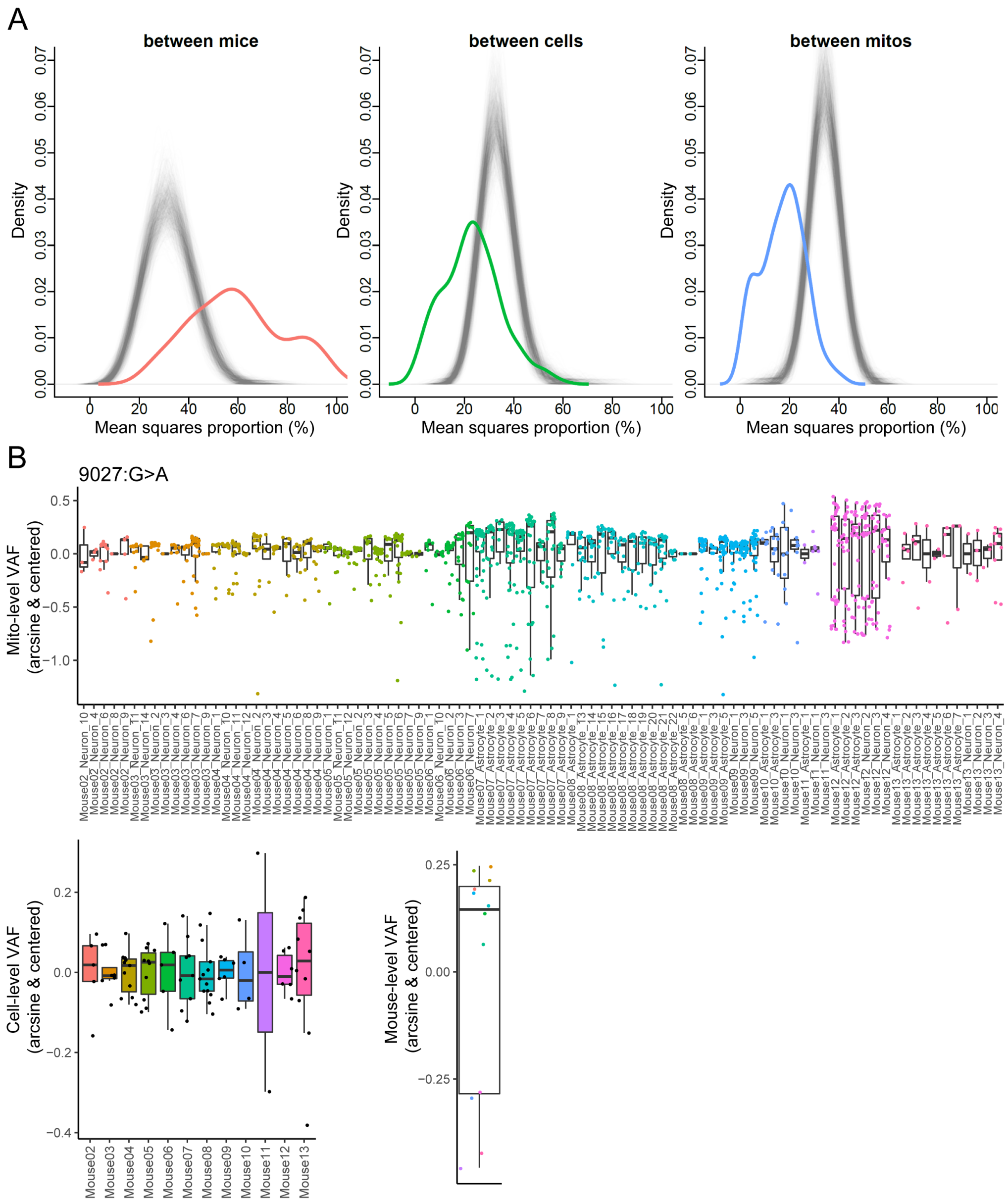


**Figure S10. Comparing inherited SNVs' AF variation across the mouse, cell, and mitochondrion level.** (A) The observed (the colored line) and the null (gray lines) distribution of the AF variation proportion at the mouse (left), cell (middle) and mitochondrion (right) level. (B) The mitochondrion-level (top), cell-level (bottom left) and mouse-level (bottom right) AF distribution of 9027: G>A. The color indicates the mouse identity.

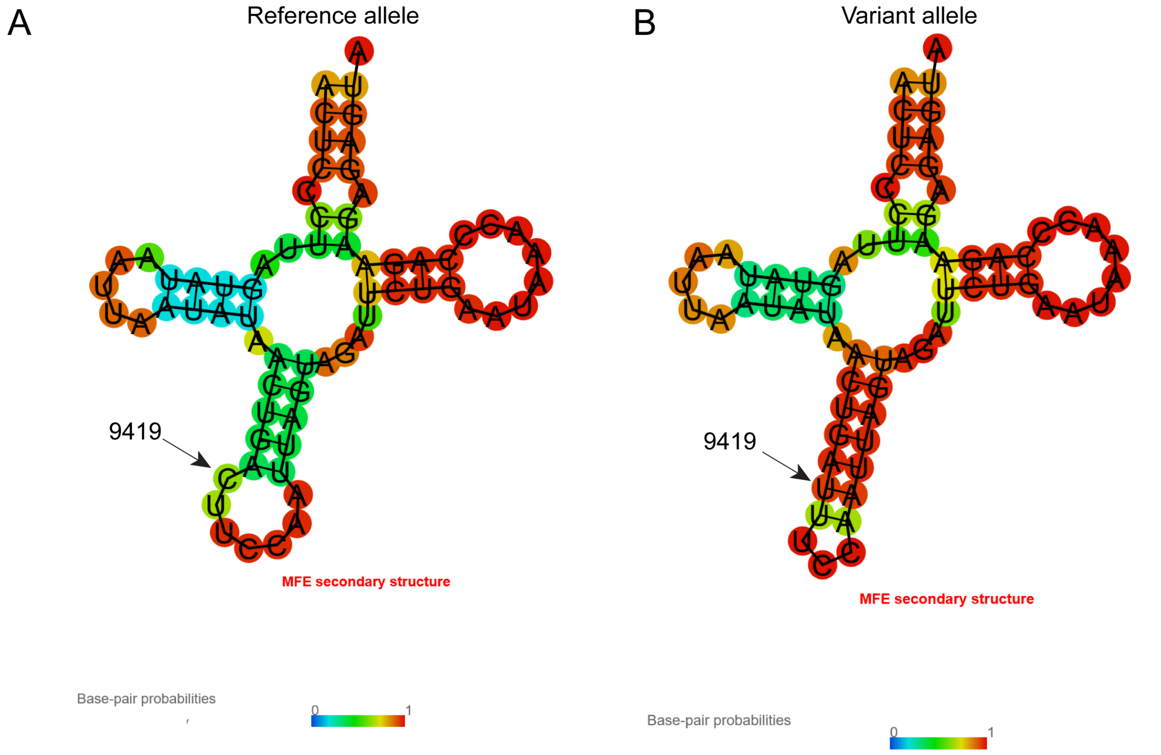


**Figure S11. RNAfold predicted tRNA secondary structure changes caused by 9419:C>T.** (A) The secondary structure of mt-Tg with the reference allele. (B) The secondary structure of mt-Tg with the variant allele.

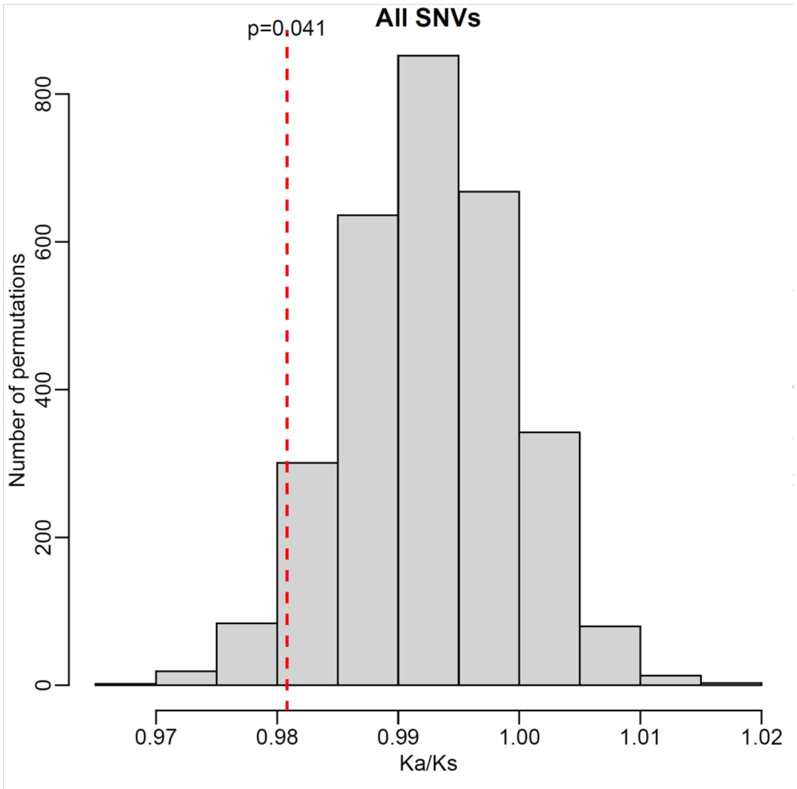


**Figure S12. Ka/Ks statistics in total SNVs.** The background distribution was derived from simulated random mutations (3000 times permutation) conditioned on the overall mutational spectrum. The red dashed-line here represents the observed Ka/Ks from total SNVs.

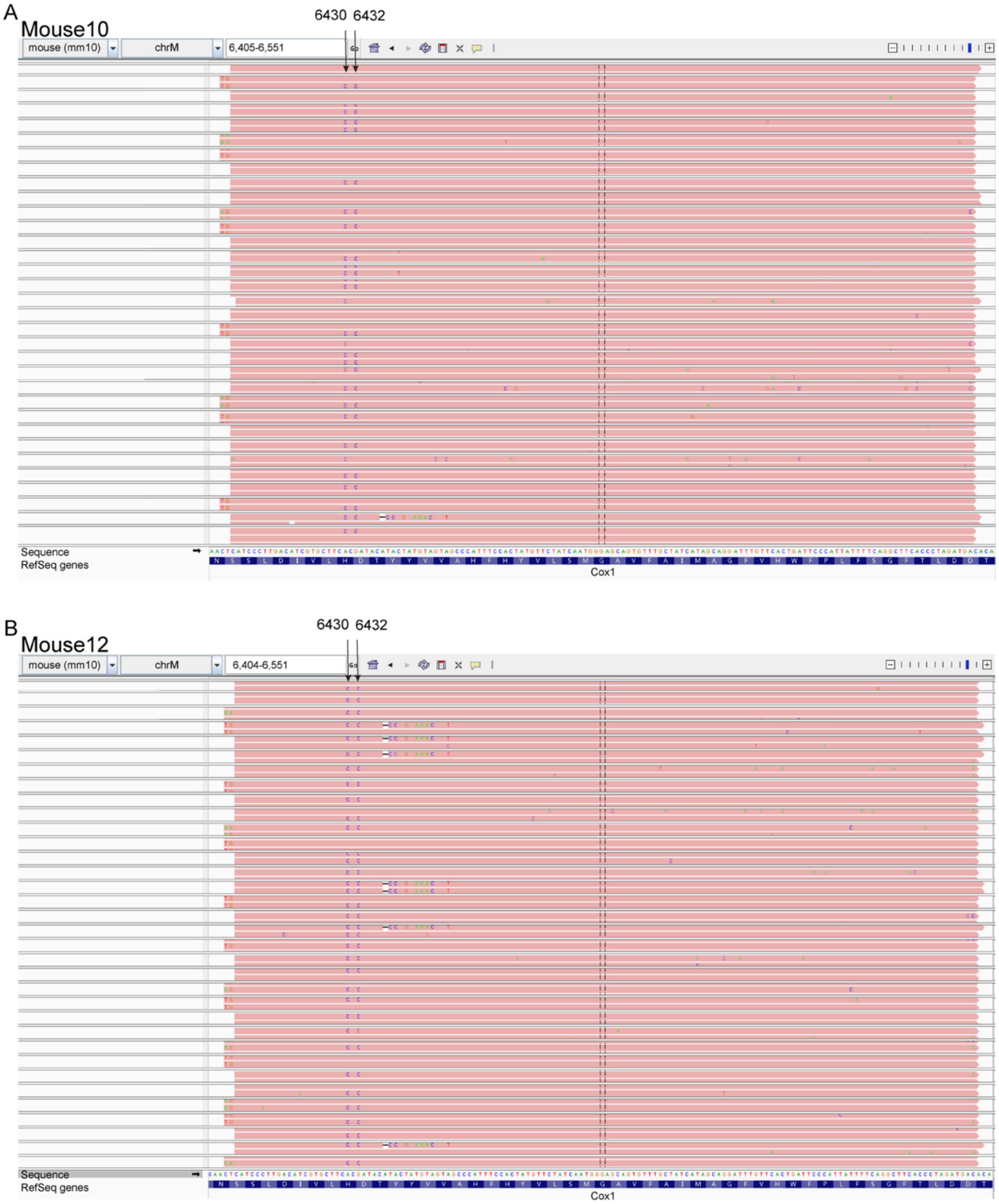


**Figure S13. IGV snapshots showing the 6430-6432 linkage on the same haplotype.**

(A) An example of mitochondrion carrying the linked SNVs in the same reads in Mouse10.

(B) An example of mitochondrion carrying the linked SNVs in the same reads in Mouse12.

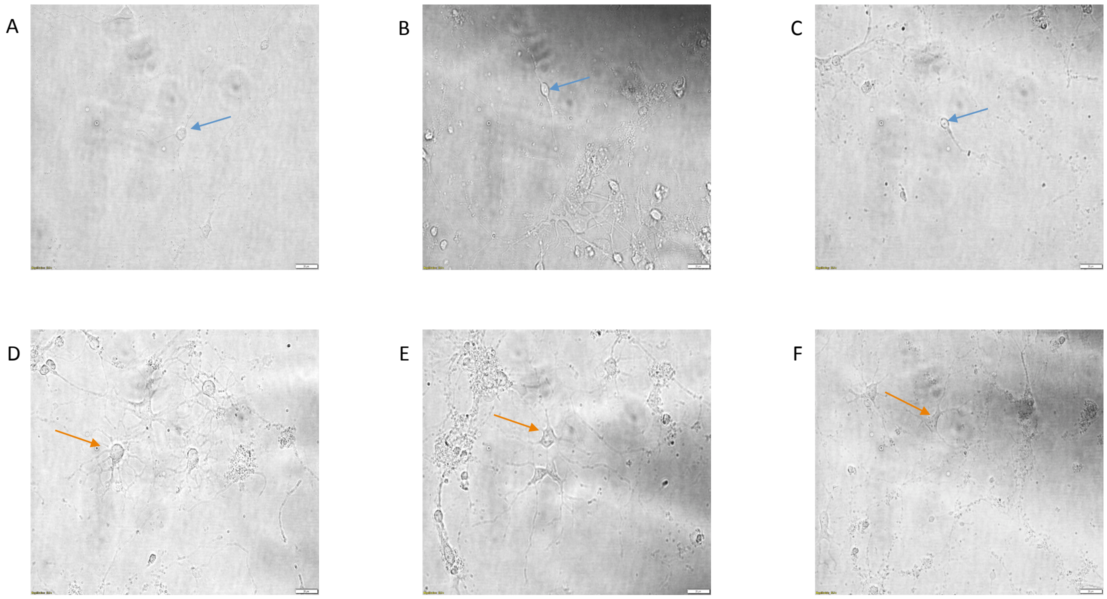


**Figure S14. Representative images of primary mouse neurons and astrocytes.** A representative image of a pyramidal (A), bipolar (B) and unipolar (C) neuron, shown by blue arrows and astrocytes in (D-F) shown by orange arrows. Scale bars are 20 microns.

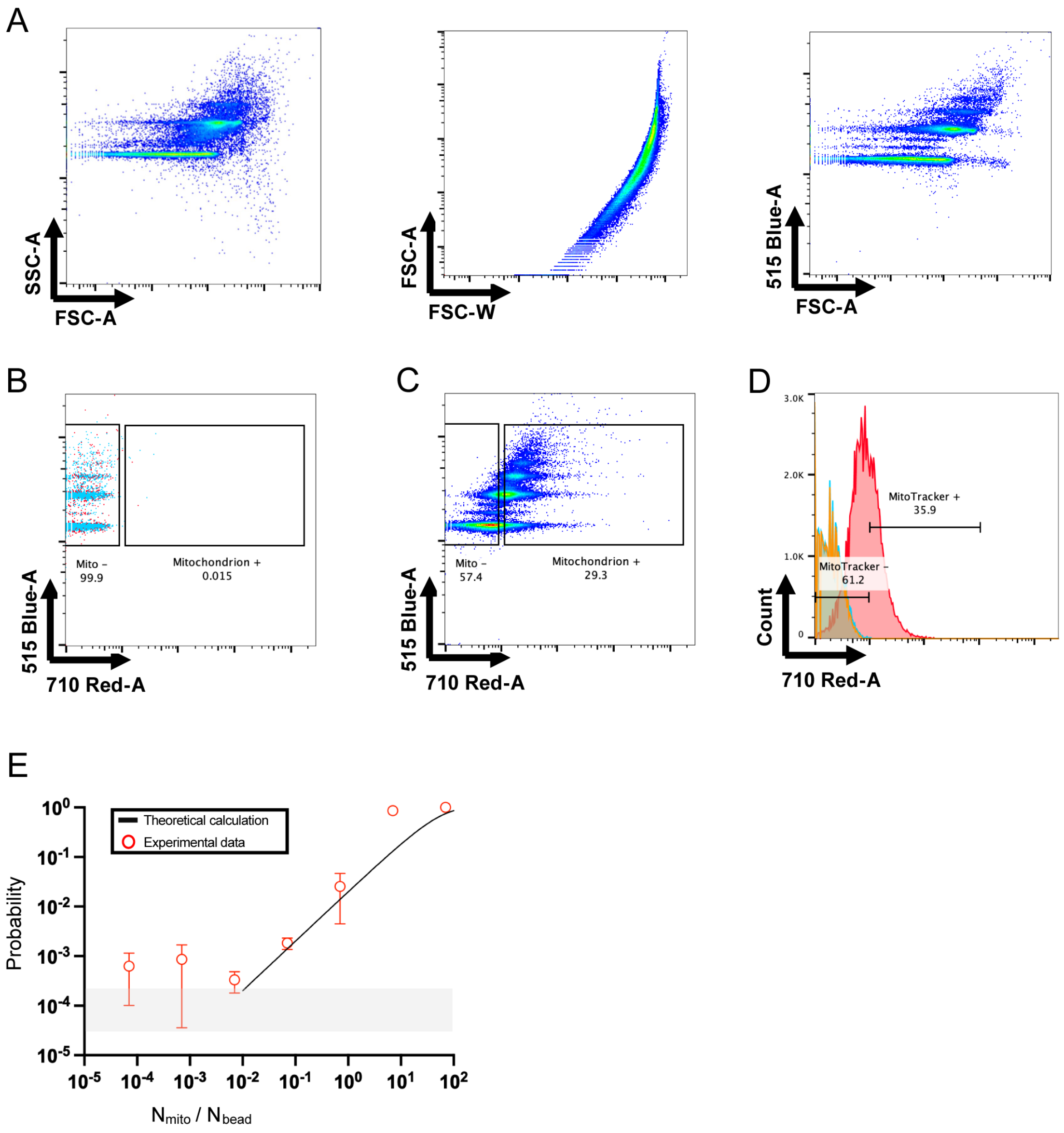


**Figure S15. Single mitochondrion isolation from individual neuron or astrocyte cells based on Poisson statistics.** (A) Flow cytometry characterization of antibody coated microbeads. (B) Gating strategy for isolating single mitochondrion from cell lysate of individual neuron or astrocyte staining by MitoTracker Red. Red dots indicated microbeads-only control, blue dots indicate microbeads captured unstained mitochondria as negative control. (C) MitoTracker gating on microbeads incubated with mitochondria stained by MitoTracker Red as positive control. (D) Comparing the MitoTracker signal from samples in B and C. Red plot indicate microbeads captured MitoTracker stained mitochondria. The orange plot depicts microbeads-only control. Blue plot depicts microbeads captured unstained mitochondria. (E) Experimental and theoretical analysis on single mt capture based on Poisson statistics. x-axis is the ratio of input mitochondria number to input microbead number. Y-axis is the probability of microbeads that captured a mitochondrion. Error bars indicate standard deviation from $n$ = 3 independent experiments.

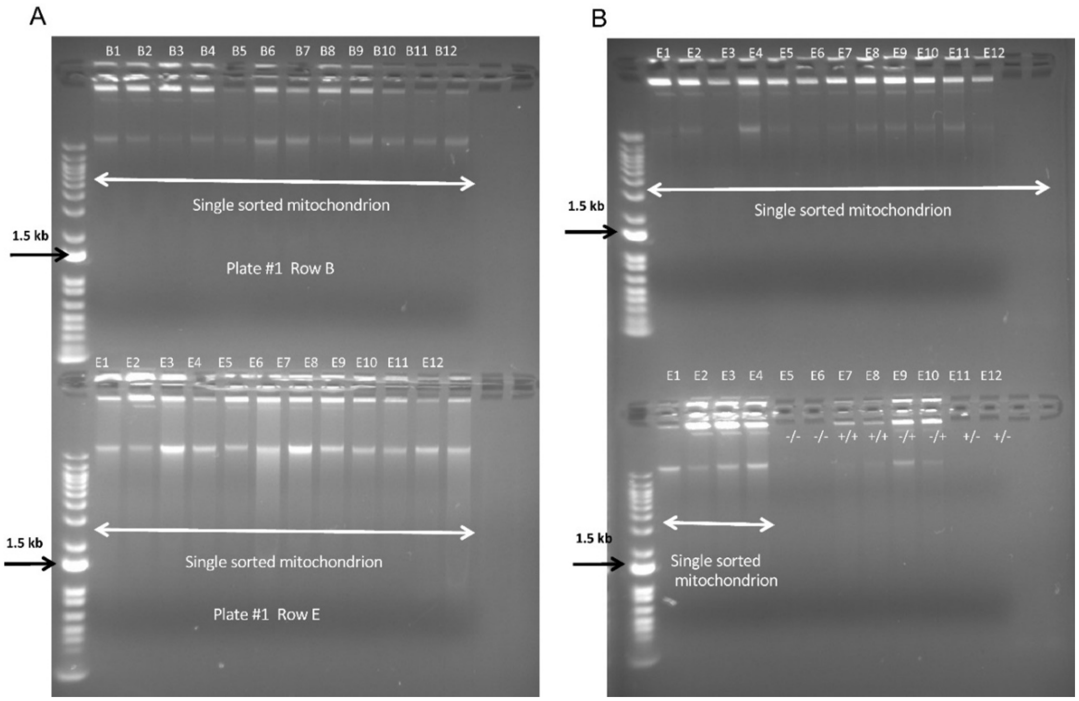


**Figure S16**. **RCA products from single mitochondrion samples.** Shown here is a representative ethidium bromide agarose gel electrophoresis with RCA products. Each well in (A) and (B) were loaded with 25-fold diluted RCA products from each mitochondrion except the control wells E5-E12 in (B). E5-6 are negative controls, E7-8 are positive controls (0.1 picogram isolated mtDNA from mouse brain used as a template), E9-10 are random primer control and E11-12 are no-template controls.

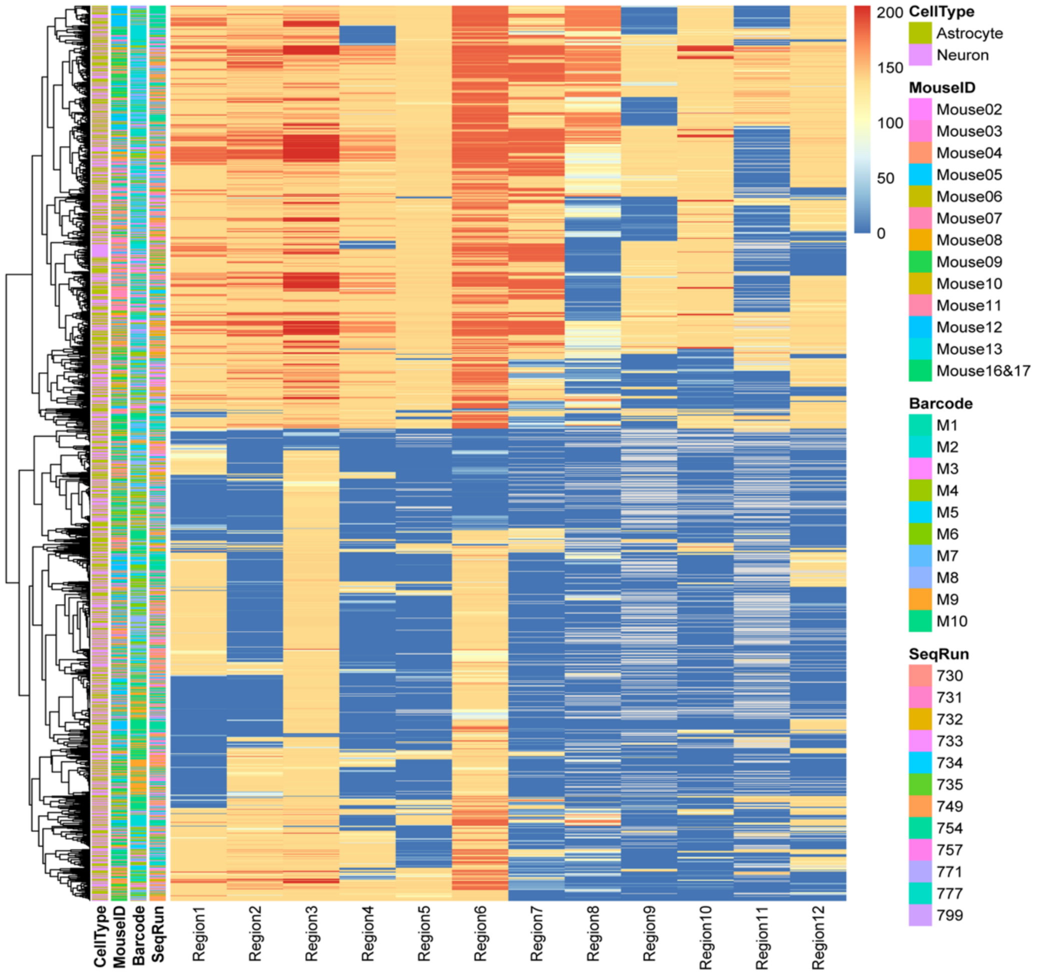


**Figure S17. Sample size and quality assessment of positive and negative controls.** The heatmap showing the number of bases with sufficient depth (≥ 50) in each mt (row) for each PCR target region (depicted as columns).

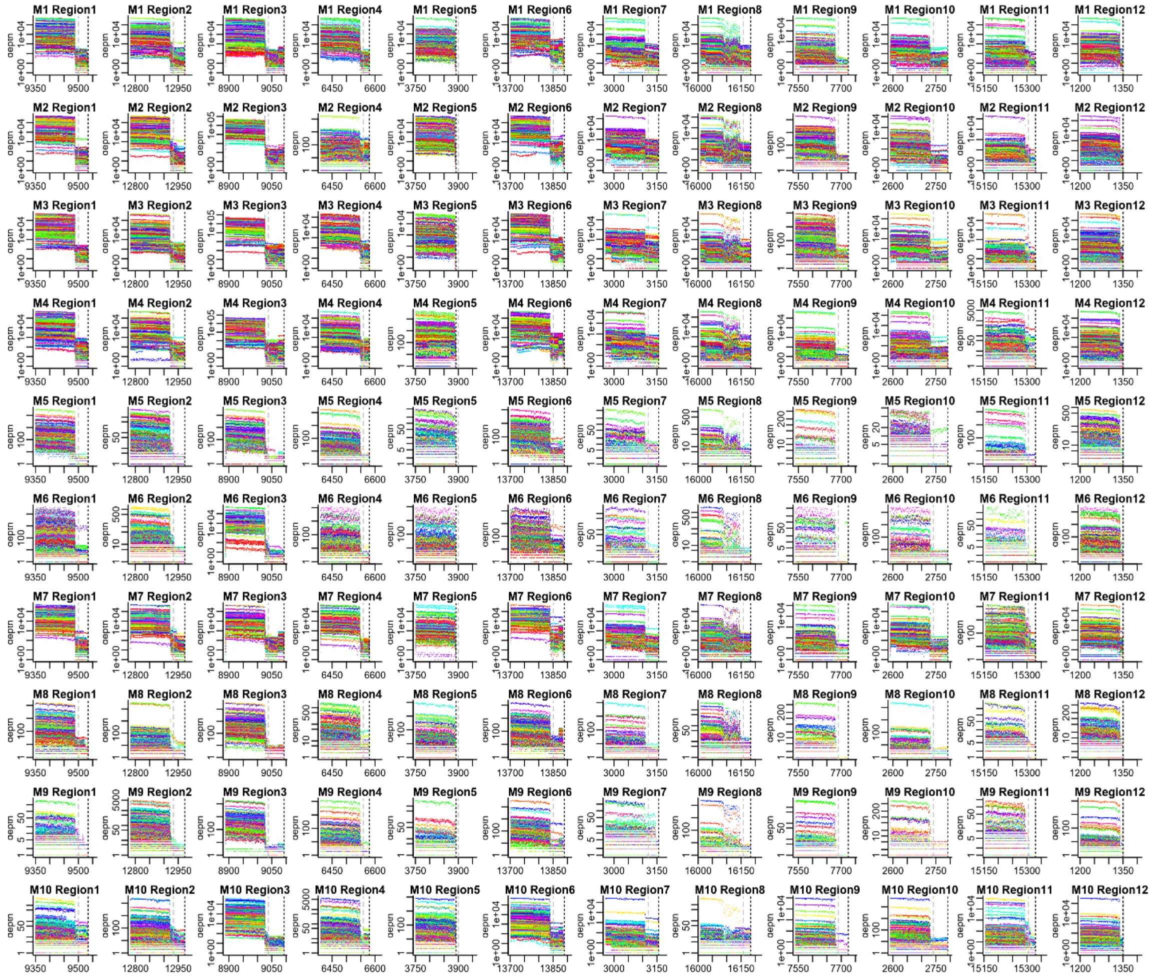


**Figure S18. Per-base read depth for each mt-barcode and each PCR target region.** Each subfigure depicts one of the 120 combinations of mt-barcode and PCR target region, and an individual curve (and color) within a subfigure represents a single mitochondrion.

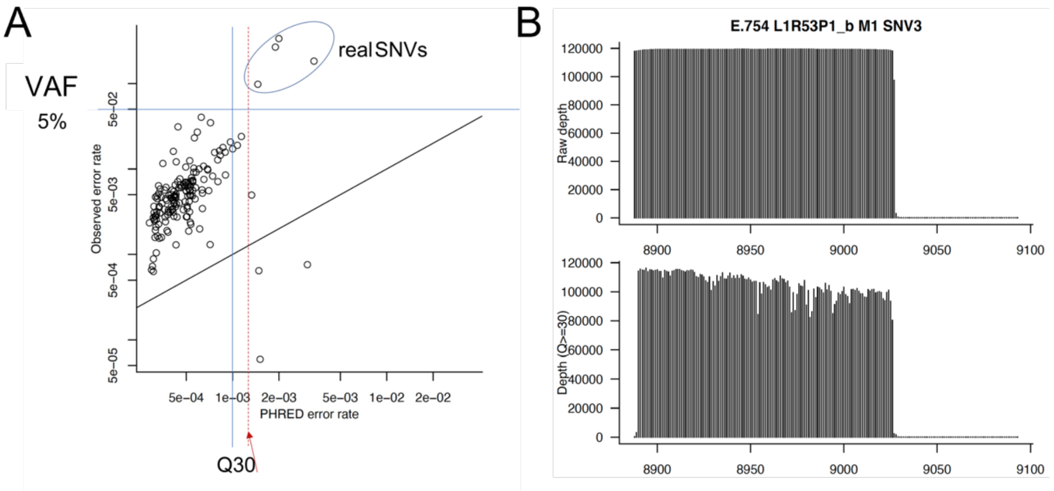


**Figure S19. Phred score threshold choice and its impact on read coverage.** (A) Scatter plot showing the theoretical error rate (Phred score, x-axis) and the observed mismatch rate (y-axis). (B) Read coverage in target Region3 before (top) and after (bottom) Q30 filtering.

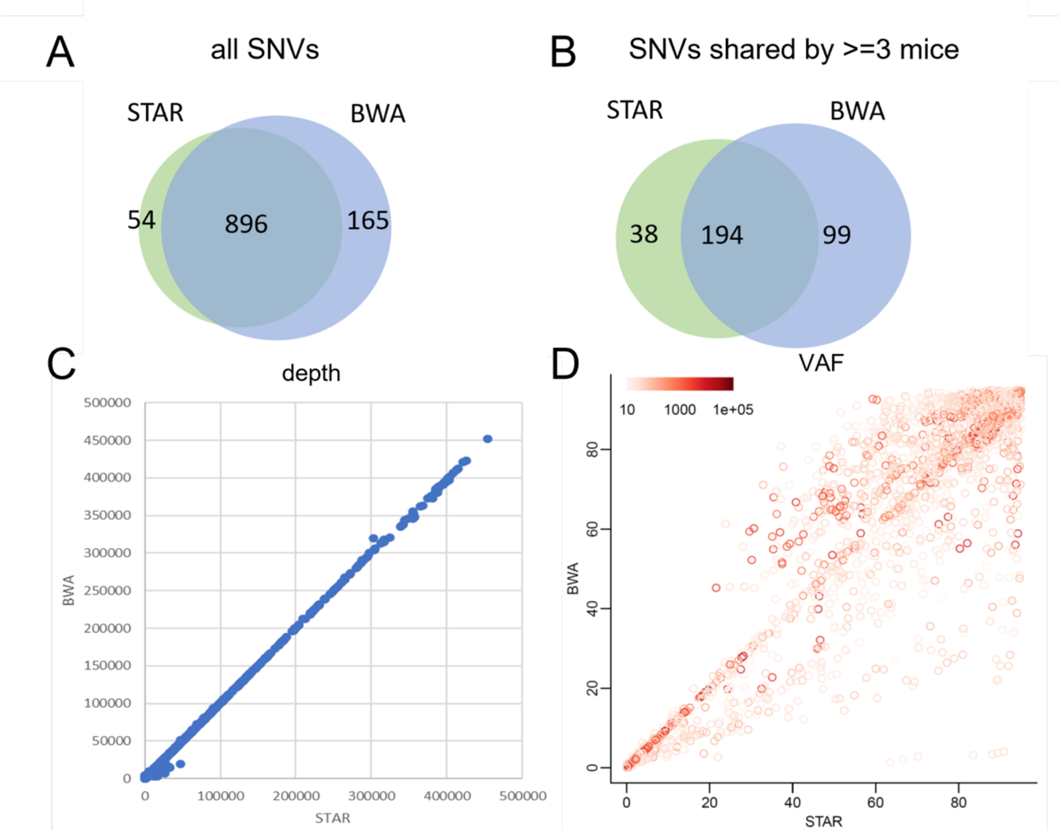


**Figure S20. STAR-BWA pipeline comparison in SNV location, depth and VAF.** (A) Venn diagram showing the overlap between STAR and BWA in all SNV sites. (B) Venn diagram showing the overlap between STAR and BWA in SNV sites shared by at least three mice. (C) Scatter plot showing the depth of STAR (x-axis) and BWA (y-axis). (D) Scatter plot showing the VAF of STAR (x-axis) and BWA (y-axis).

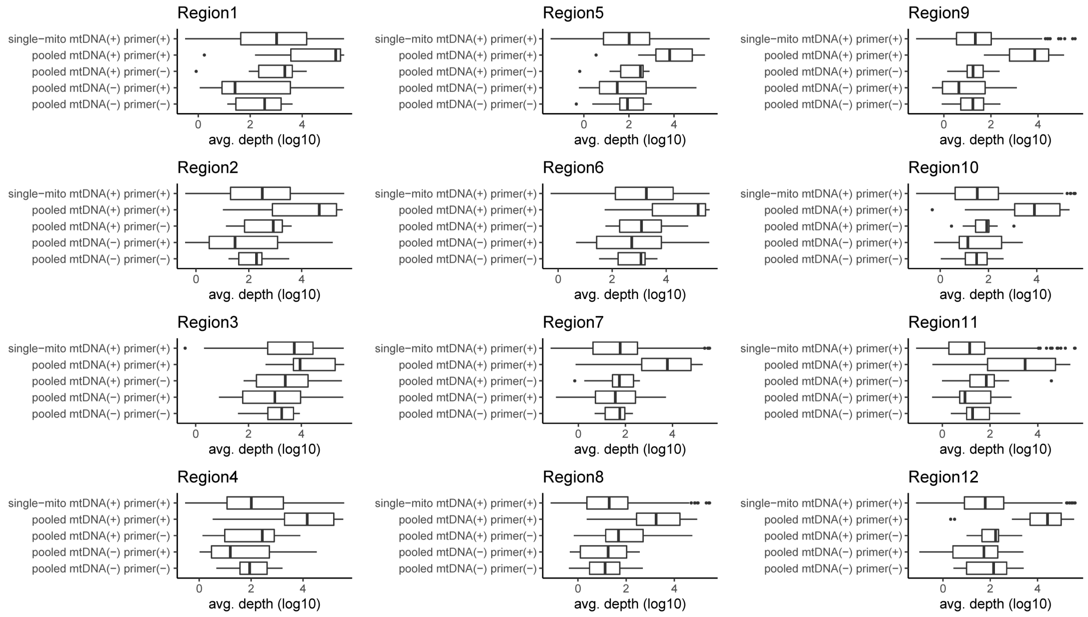


**Figure S21. Boxplots of the read depth in each of the 12 PCR target regions. The average read depth (log 10)** for single-mt samples [single-mtDNA (+) primer (+)] and positive [pooled mtDNA (+) primer (+)] or negative controls [mtDNA (-) RCA (-)] or other controls [mtDNA (+) primer (-), mtDNA (-) primer (+)] in Region 1 – 12.

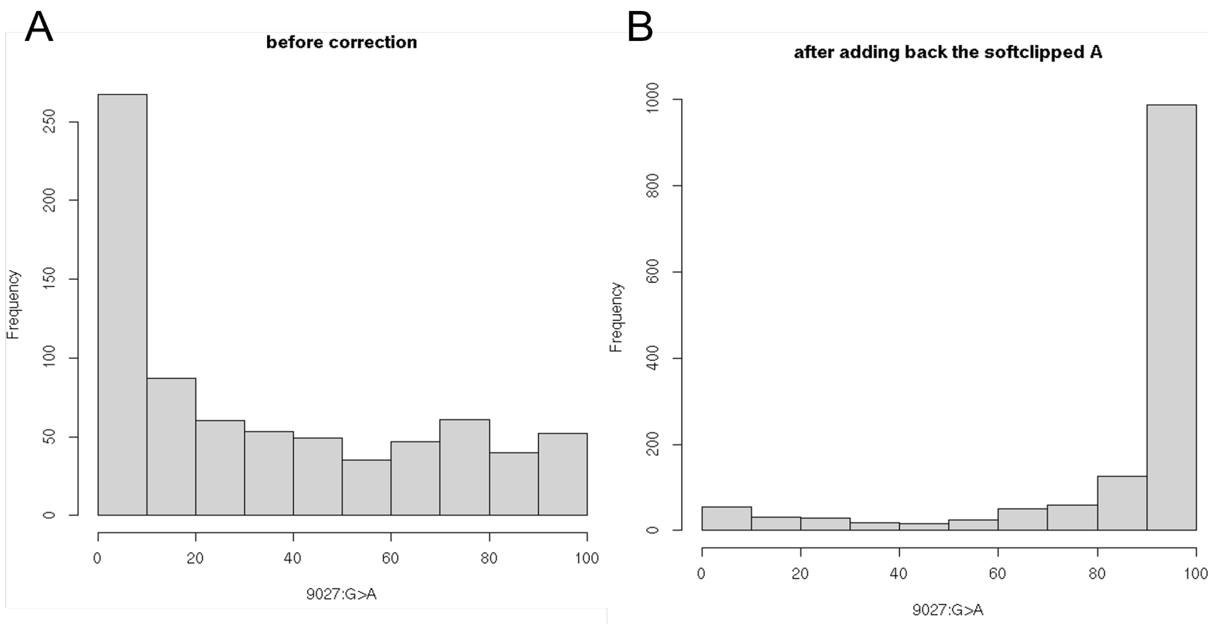


**Figure S22. Correction for Soft-clipping by STAR on 9027:G>A.** The VAF difference between before and after patching the non-reference base As that were soft-clipped by STAR. (A) Before the correction. (B) After the correction.

A

B

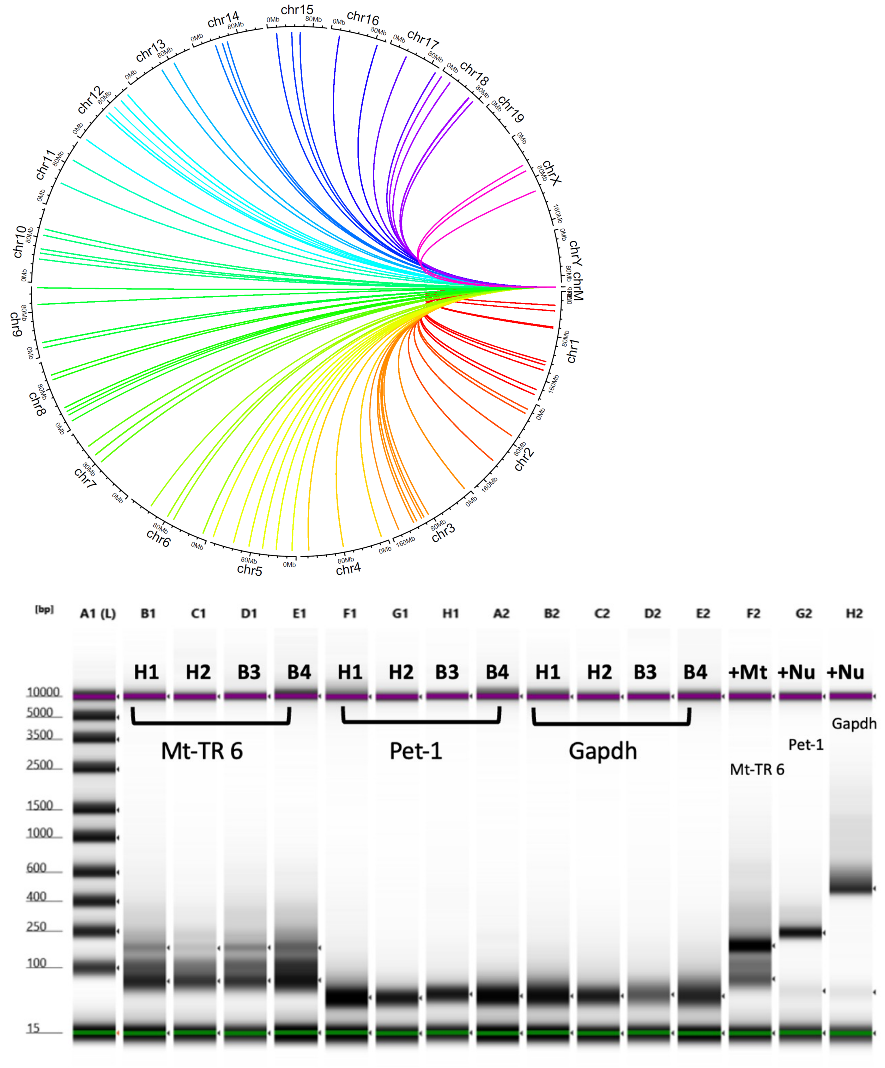


**Figure S23. Mouse NUMTs Analysis on the SMITO dataset.** (A) Homologous mtDNA shown as links; colors denote individual chromosomes. (B) Representative example of D5000 Agilent ScreenTape with PCR product from targeting single mt RCA product for a mt region (SNV6) and 2 independent nuclear regions (Pet-1 and Gapdh).

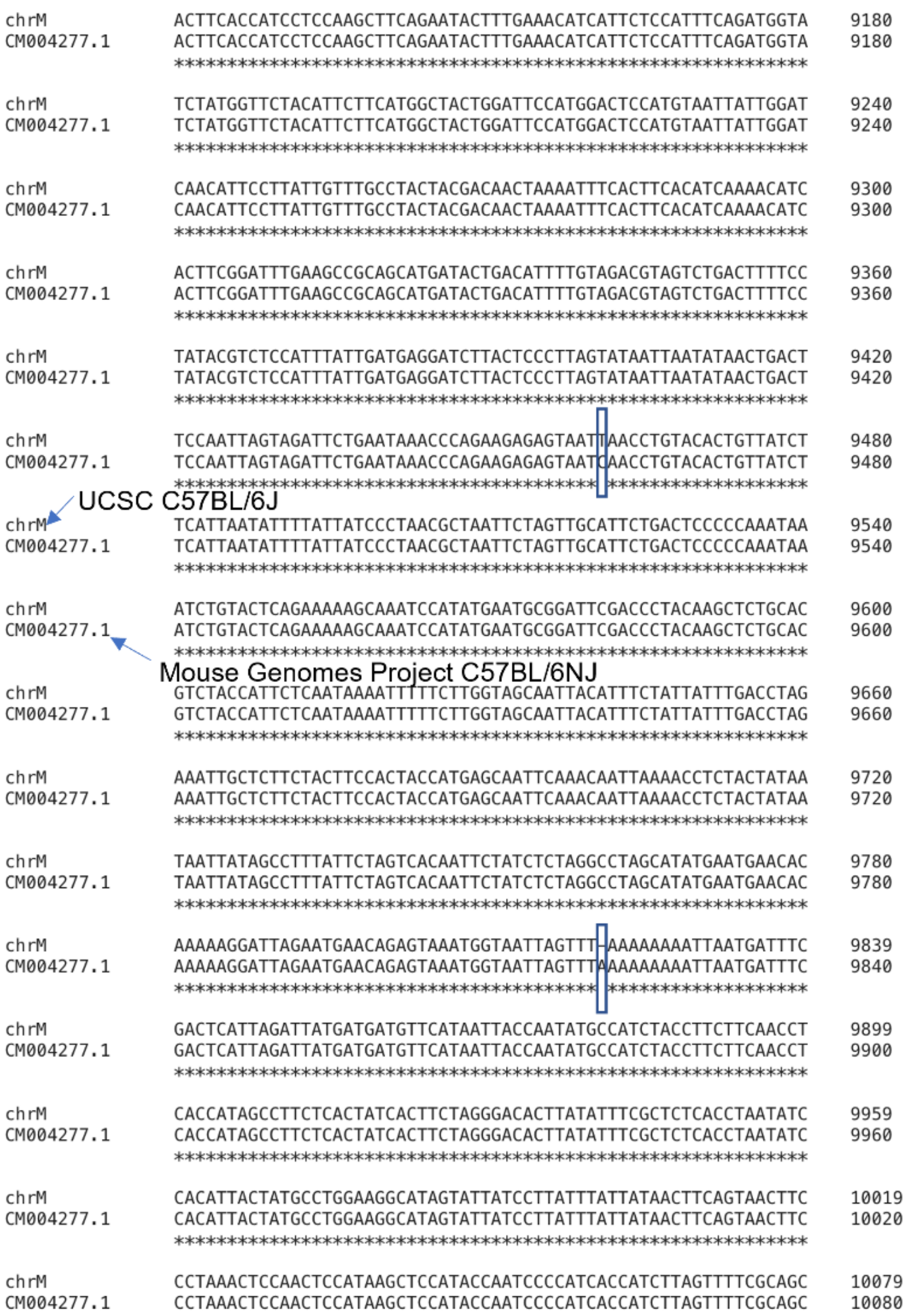


**Figure S24. Sequence comparison between UCSC GRCm38 (C57BL/6J) and C57BL/6NJ from the Mouse Genomes Project in the region highlighting two differences.**

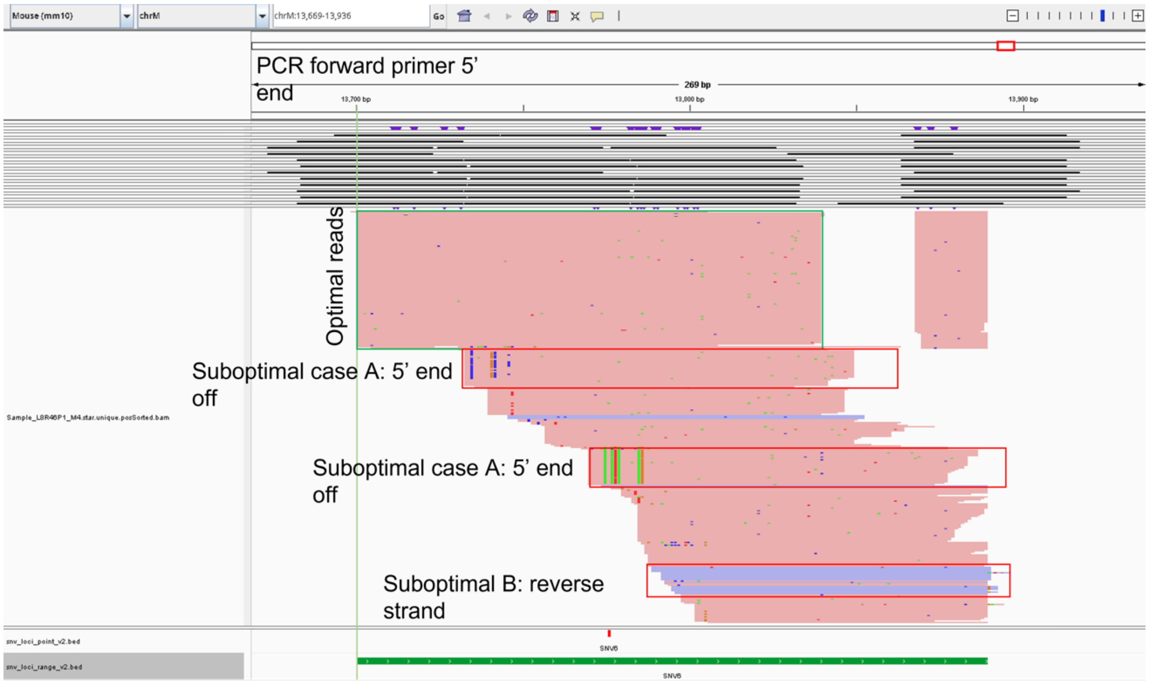


**FigureS25. Examples of suboptimal alignments and the mutations in linkage.** The green box at the top highlights the optimal alignments, while the three red boxes represent suboptimal cases.

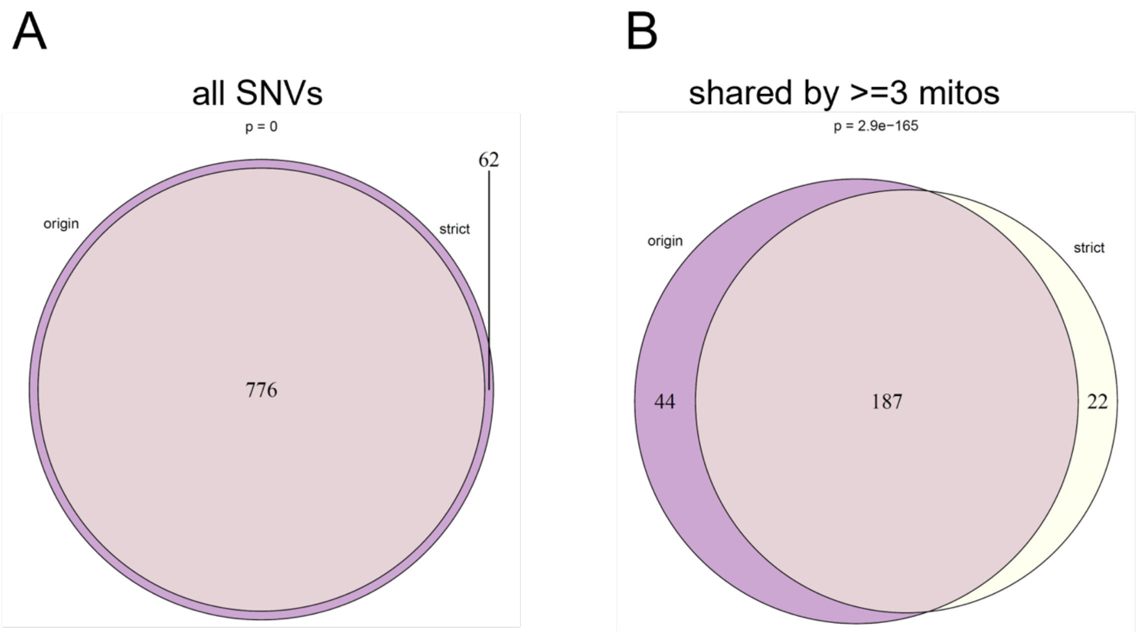


**Figure S26. Comparison between the permissive and the stricter filtering.** (A) Venn diagram showing the overlap of all SNV sites between the permissive and the stricter filtering. (B) Venn diagram showing the overlap of the SNV sites shared by at least 3 mitochondria between the permissive and the stricter filtering.

**Supplementary Tables.**

**Table S1A**. List of the sequences of the 10 barcodes (M1-10) for mitochondria multiplexing.

| ID | Spacer | Barcode | Full sequence |
|---|---|---|---|
| M1 | CGATT | ATCACG | CGATTATCACG |
| M2 | CGTAT | CGATGT | CGTATCGATGT |
| M3 | GCTAA | TTAGGC | GCTAATTAGGC |
| M4 | GCTTA | TGACCA | GCTTATGACCA |
| M5 | CGATT | ACAGTG | CGATTACAGTG |
| M6 | CGTAT | GCCAAT | CGTATGCCAAT |
| M7 | CGTAT | CAGATC | CGTATCAGATC |
| M8 | CGTAT | GATCAG | CGTATGATCAG |
| M9 | CGTAT | AGTCAA | CGTATAGTCAA |
| M10 | CGTAT | ACTGAT | CGTATACTGAT |

**Table S1B.** Edit distances between any pair of the mitochondrial-barcodes.

| Levenshtein | M1 | M2 | M3 | M4 | M5 | M6 | M7 | M8 | M9 | M10 |
|---|---|---|---|---|---|---|---|---|---|---|
| M1 | 0 | 5 | 6 | 5 | 3 | 6 | 6 | 3 | 5 | 6 |
| M2 | 5 | 0 | 7 | 6 | 6 | 4 | 3 | 4 | 5 | 4 |
| M3 | 6 | 7 | 0 | 5 | 7 | 9 | 6 | 7 | 7 | 7 |
| M4 | 5 | 6 | 5 | 0 | 7 | 5 | 6 | 4 | 5 | 6 |
| M5 | 3 | 6 | 7 | 7 | 0 | 5 | 5 | 5 | 6 | 5 |
| M6 | 6 | 4 | 9 | 5 | 5 | 0 | 4 | 4 | 3 | 3 |
| M7 | 6 | 3 | 6 | 6 | 5 | 4 | 0 | 4 | 4 | 3 |
| M8 | 3 | 4 | 7 | 4 | 5 | 4 | 4 | 0 | 3 | 4 |
| M9 | 5 | 5 | 7 | 5 | 6 | 3 | 4 | 3 | 0 | 3 |
| M10 | 6 | 4 | 7 | 6 | 5 | 3 | 3 | 4 | 3 | 0 |

**Table S2.** List of the sequences of the forward/reverse PCR primers for selected specific target regions.

| Target region | SNV position | Gene | Forward primer sequence 5' to 3' | Reverse primer sequence 5' to 3' |
|---|---|---|---|---|
| 1 | 9461 | mt-Nd3 | CTGACTTTTCCTATACGTCTCCA | GGGGGAGTCAGAATGCAACTA |
| 2 | 12913 | mt-Nd5 | CATAGCCTGGCAGACGAACA | ATTAGTAGGGCTCAGGCGTTG |
| 3 | 9027 | mt-Co3 | TGCAGGATTCTTCTGAGCGT | GGGCTTGATTTATGTGGTTTCGT |
| 4 | 6543 | mt-Co1 | CATCCCTTGACATCGTGCTTC | AATATGATGGCGAAGTGGGCT |
| 5 | 3816 | mt-Tq | AGAGGTTCAAGCCCTCTTATTT | CAACGTTTTCGGGGTATGGG |
| 6 | 13776 | mt-Nd6 | ACCAATCTCCCAAACCATCAAG | GGGGGATGTTGGTTGTGTTT |
| 7 | 3079 | mt-Nd1 | GCACCTACCCTATCACTCACAC | CGGCTCGTAAAGCTCCGAA |
| 8 | 16029 | D-Loop | GTCCGCAAAACCCAATCACC | TGATCAGGACATAGGGTTTGATAGT |
| 9 | 7612 | mt-Co2 | AGGCCGACTAAATCAAGCAA | AGGTTAACGCTCTTAGCTTC |
| 10 | 2651 | mt-Rnr2 | ACCTTACAAATAAGCGCTCTCAAC | TAGAATGGGGACGAGGAGTGT |
| 11 | 15191 | mt-Cytb | AATTGGGGGCCAACCAGTAG | TTCAGGTTTACAAGACCAGAGT |
| 12 | 1317 | mt-Rnr2 | ATAGAACTAGTACCGCAAGGGA | GTAGCTCGTTTGGTTTCGGG |

**<u>Note: The following supplementary datasets are provided as individual excel files.</u>**

**Legends for Datasets:**

**Dataset S1 (separate file).** The metadata for each mitochondrion or control sample.

**Dataset S2 (separate file).** The impact annotation for all 1032 SNVs with SIFT scores.

**Dataset S3 (separate file).** Major alleles and top minor alleles of SMITO and the 17 mouse strains data.

**Dataset S4 (separate file).** Ti/Tv comparison (genera, species, strains, and populations).

**Dataset S5 (separate file).** MK test (castaneus-vs-domesticus).

**Dataset S6 (separate file).** Differential statistics from the cell-type comparison in SNV incidence and AF variance.

**Dataset S7 (separate file).** Ka/Ks statistic for astrocytes and neurons.

**Dataset S8 (separate file).** The SNV AF and the number of sample support table.

**Dataset S9 (separate file).** Quality check statistics from pipeline pre-processing.

**Dataset S10 (separate file).** Mouse NUMTs analysis on SMITO data.
